# Enhancer-driven gene regulatory networks inference from single-cell RNA-seq and ATAC-seq data

**DOI:** 10.1101/2022.12.15.520582

**Authors:** Yang Li, Anjun Ma, Yizhong Wang, Qi Guo, Cankun Wang, Shuo Chen, Hongjun Fu, Bingqiang Liu, Qin Ma

**Author notes:** To whom correspondence should be addressed, BL, QM. Email addresses: YL AM YW QG CW SC HF.

## Abstract

Deciphering the intricate relationships between transcription factors (TFs), enhancers, and genes through the inference of enhancer-driven gene regulatory networks is crucial in understanding gene regulatory programs in a complex biological system. This study introduces STREAM, a novel method that leverages a Steiner Forest Problem model, a hybrid biclustering pipeline, and submodular optimization to infer enhancer-driven gene regulatory networks from jointly profiled single-cell transcriptome and chromatin accessibility data. Compared to existing methods, STREAM demonstrates enhanced performance in terms of TF recovery, TF-enhancer relation prediction, and enhancer-gene discovery. Application of STREAM to an Alzheimer’s disease dataset and a diffuse small lymphocytic lymphoma dataset reveals its ability to identify TF-enhancer-gene relationships associated with pseudotime, as well as key TF-enhancer-gene relationships and TF cooperation underlying tumor cells.

## BACKGROUND

Recent advancements in single-cell sequencing technologies, such as RNA sequencing (scRNA-seq) and Assay for Transposase-Accessible Chromatin using sequencing (scATAC-seq), have greatly contributed to the elucidation of gene regulatory networks (GRNs) at a single-cell resolution^1^. Current transcriptome-based methods for constructing GRNs combine gene co-expression analysis with discovering transcription factor (TF) binding motifs. For example, SCENIC, a random forest-based computational method tailored to deduce GRNs and pinpoint cell types or states from scRNA-seq data^2^. DIRECT-NET adopts frequentist non-linear regression to spotlight *cis*-regulatory elements (CREs) and GRNs^3^. scMEGA harnesses trajectories to infer GRNs linearly^4^. However, this motif discovery process focuses on searching for TF binding sites within a limited sequence space surrounding each co-expressed gene module, thereby only capturing a fraction of a gene’s potential regulatory space. Moreover, traditional motif finding approaches primarily base their predictions of TF binding sites on sequence conservation, and they often overlook the variability in chromatin accessibility and regulatory sequences, which can result in high false-positive rates in TF predictions when applied to single-cell data^5^. Such limitations induced by a single modality profiling can be overcomed by the integration of single-cell multi-omics data. GRN inference through integrative analysis of scRNA-seq and scATAC-seq data can reduce the impact of noise from sparse sequencing and improves the accuracy of TF-target predictions by validating regulatory relationships across different modalities^1,3,4,6-19^. Motif discovery is no longer restricted to limited promoter regions, and the heterogeneity of regulatory sequences can be preserved. On top of the advances, adding chromatin accessibility information into GRN enables the inference of enhancer-driven GRNs (eGRNs)^7^. In an eGRN, the edges connecting TFs to target genes pass through binding enhancers^7^, where enhancers refer to regions on chromatins that participate in gene regulation via TF binding.

The elucidation of eGRNs offers new insights to the inference of cell-type-specific/conserved TF regulatory patterns and the variability of TF-enhancer and enhancer-gene relations^20,21^. Recently, Janssens, *et al*., introduced the concept of eGRNs and developed an extensive resource of eGRNs for 40 cell types in the fly brain using a bioinformatic framework equipped with cell clustering, motif discovery, network prediction, and deep learning models^7^. Central to this advancement is SCENIC+^18^, which is a cutting-edge tool adept at deducing eGRNs, integrating SCENIC and pycisTopic frameworks. It stands out by predicting genomic enhancers, identifying associated upstream TFs, and linking these enhancers to their potential target genes. To reinforce the precision of TF identification, SCENIC+ incorporates a meticulously curated motif collection of over 30,000 motifs. Its versatility is evident as it has been benchmarked across diverse datasets from various species, covering human peripheral blood mononuclear cells, ENCODE cell lines, to specific cases like melanoma states and Drosophila retinal development. Another tool, GLUE, employs a deep learning approach to discern cell clusters and deduce GRNs from single-cell data spanning transcriptomes, chromatin accessibility, and DNA methylation^6^. Lastly, Pando shapes GRNs by capitalizing on scRNA-seq and scATAC-seq, modeling gene expression via transcription factor-peak interactions^22^.

Presently, the inference of eGRNs poses three key challenges for method development. First, existing approaches for predicting enhancer-gene relationships rely on profile correlations between enhancer accessibility and gene expression. For example, SCENIC+^18^, SCENIC^18^, GLUE^6^, DIRECT-NET^3^, Pando^16^, and scMEGA^4^ predict TF-enhancer-gene or TF-gene relations by treating each relation individually rather than considering their mutual dependency. Hence, these methods often suffer from a high false positive rate in complex scenarios where multiple enhancer-enhancer and enhancer-gene relationships cooperatively participate in gene regulation. Second, current methodologies typically identify co-expressed or co-regulated genes and their associated enhancers in a cell type by leveraging prior cell clustering in a sequential manner. Biases introduced during the preceding clustering step can subsequently affect the accuracy of predicting TF-enhancer-gene relationships. Last, the regulatory heterogeneity present in cells gives rise to numerous combinations of TF-enhancer-gene relationships that govern cell states and behaviors. Unraveling the global extraction of key TF-enhancer-gene relationships across all cells becomes intricate due to this regulatory heterogeneity.

To address these challenges, we propose STREAM (<u>S</u>ingle-cell enhancer regula<u>T</u>ory netwo<u>R</u>k inference from gene <u>E</u>xpression <u>A</u>nd chro<u>M</u>atin accessibility), a computational framework for eGRN inference from jointly profiled scRNA-seq and scATAC-seq data (**Fig. 1a**) using Steiner Forest problem (SFP) model and submodular optimization. STREAM constructs a heterogeneous graph to represent enhancer-enhancer and enhancer-gene relationships. Utilizing the SFP model, it identifies robust enhancer-gene associations that constitute a context-specific functional gene module (FGM)^23^. In its eGRN inference, STREAM detects hybrid biclusters (HBCs), highlighting genes co-regulated by a common TF and associated co-accessible enhancers. This approach obviates the need for prior cell clustering. Subsequently, through submodular optimization, STREAM selects the most representative combination of HBCs, emphasizing primary TF-enhancer-gene interactions^24^. As a result, the TF, enhancers, and genes found within the same HBC, along with their interrelationships, together constitute an enhancer-driven Regulon (eRegulon). Subsequently, eRegulons active within a specific cell type form the basis of an eGRN.

**Fig. 1.**
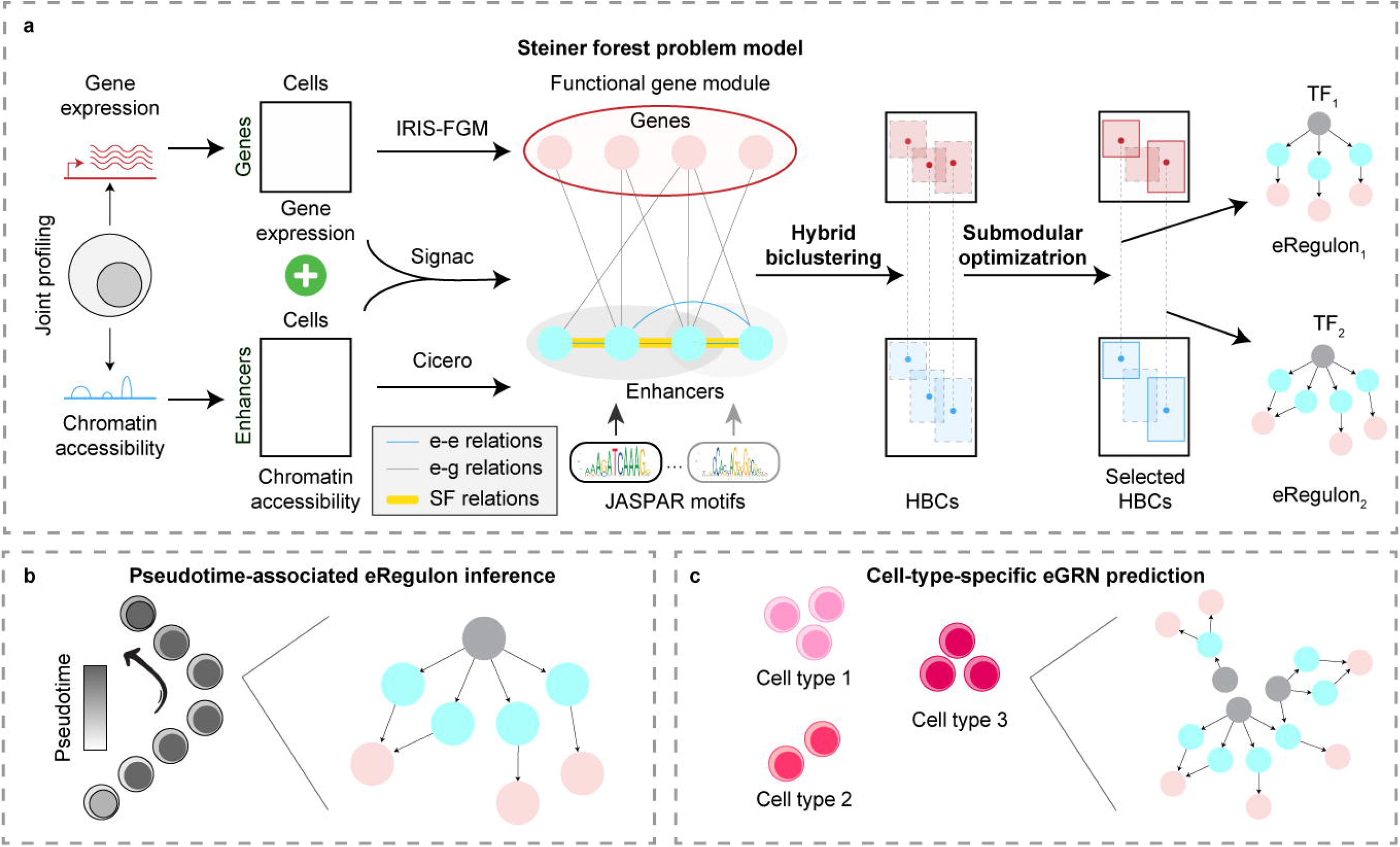
STREAM Method Overview and Applications in AD and DSLL. **a**. An outline of the STREAM framework for eRegulon identification. An eRegulon comprises a TF, its set of binding enhancers, and target genes. **b**. STREAM’s application in predicting pseudotime-associated eRegulons in AD. **c**. The use of STREAM for inferring cell-type-specific eGRNs in DSLL.

To evaluate the performance of STREAM, we applied it to six jointly profiled scRNA-seq and scATAC-seq datasets generated using various technologies and compared the results with SCENIC+^18^, SCENIC^2^, GLUE^6^, DIRECT-NET^3^, Pando^22^, and scMEGA^4^. The evaluation criteria in benchmarking analyses include TF recovery, TF-enhancer relation prediction, and enhancer-gene relation discovery. The comparison results revealed that STREAM performed better than these methods for (e)GRN inference. We then applied STREAM to an Alzheimer’s disease (AD) dataset over a time course and unveiled pseudotime-associated eRegulons and changing enhancer-gene relations alongside cell lineages (**Fig. 1b**). Additionally, by running STREAM on a diffuse small lymphocytic lymphoma (DSLL) dataset, we demonstrated its capacity to identify eRegulons underlying the diseased cell type and showcase the cooperation among TFs in gene regulation (**Fig. 1c**). These findings highlight the competing performance and versatility of STREAM in inferring eGRNs from jointly profiled scRNA-seq and scATAC-seq data, and provide valuable insights into gene regulatory mechanisms underlying complex biological processes and diseases. The STREAM tool is available as an R package on GitHub (https://github.com/YangLi-Bio/stream_core), offering accessibility for further exploration.

## RESULTS

### Overview of the STREAM pipeline

The STREAM computational framework (**Fig. 2** and and **Supplementary Fig. 1**) is a sophisticated approach to eGRN inference from jointly profiled scRNA-seq and scATAC-seq data. It encompasses three integral components, each playing a pivotal role in enhancing the accuracy and depth of regulatory network identification. First, as highlighted in **Fig. 2a**, STREAM uses the SFP model, which acts as a cornerstone in the entire framework. Through the SFP model, STREAM builds a heterogeneous graph that portrays the coordinated relationships among enhancer-enhancer and enhancer-gene relations. This graph is fundamental as it facilitates the identification of robust enhancer-gene relationships, forming the foundation of a context-specific FGM. The second component, delineated in **Fig. 2b**, involves a hybrid biclustering algorithm. This algorithm is specifically designed to uncover genes that are co-regulated by the same TF and share co-accessible enhancers. An intrinsic advantage of STREAM’s approach here is its ability to discern TF-enhancer-gene relations and the related cell subpopulation of a HBC without relying on prior cell clustering. Finally, as seen in **Fig. 2c**, STREAM employs submodular optimization, a technique tailored to extract an optimal combination of HBCs, which essentially comprises the key TF-enhancer-gene relations on a global scale. By doing so, it curates a set of pivotal eRegulons. These eRegulons, when active in a particular cell type or subpopulation, together form the overarching eGRN. By meticulously integrating these functionalities, STREAM stands out as a formidable tool for architecting an eGRN. This architecture is achieved by seamlessly interconnecting the active eRegulons within a defined cell subpopulation. Further details and a granular breakdown of the STREAM pipeline are elaborated upon in the **Methods** section.

**Fig. 2.**
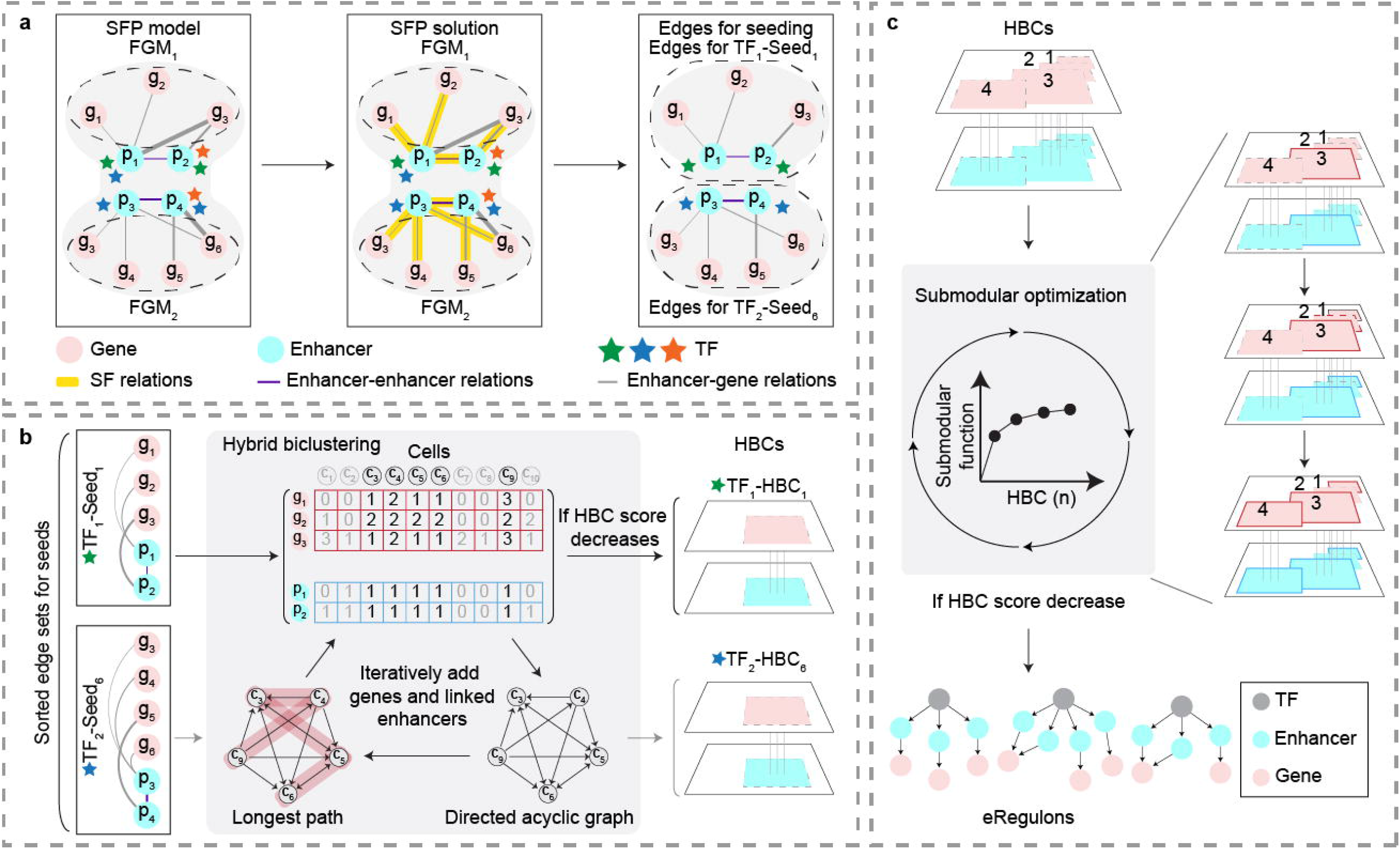
Detailed Overview of the STREAM Framework. **a**. The SFP model is utilized to extract highly confident enhancer-gene relations. **b**. The hybrid biclustering pipeline is used for the identification of HBCs. **c**. The construction of eRegulons using a submodular optimization approach based on HBCs.

### Benchmark evaluations of STREAM

We performed benchmarking analyses on six single-cell RNA-seq and ATAC-seq datasets sourced from 10X Genomics Multiome, scCAT-seq^25^, SHARE-seq^26^, and SNARE-seq2^27^ (**Fig. 3a**). These datasets were derived from cell lines including bone marrow, K562, HCT116, Hela-S3, GM12878, and A549. Our analyses sought to validate STREAM’s predictions related to TFs, their associated binding enhancers, and the relationships between enhancers and genes. While we adopted the benchmarking approach from SCENIC+^18^, our major distinction was the direct use of real biological datasets rather than simulated ones. This decision stemmed from our belief that real data provides deeper insights into the efficacy of methods and nuances of different sequencing techniques.

**Fig. 3.**
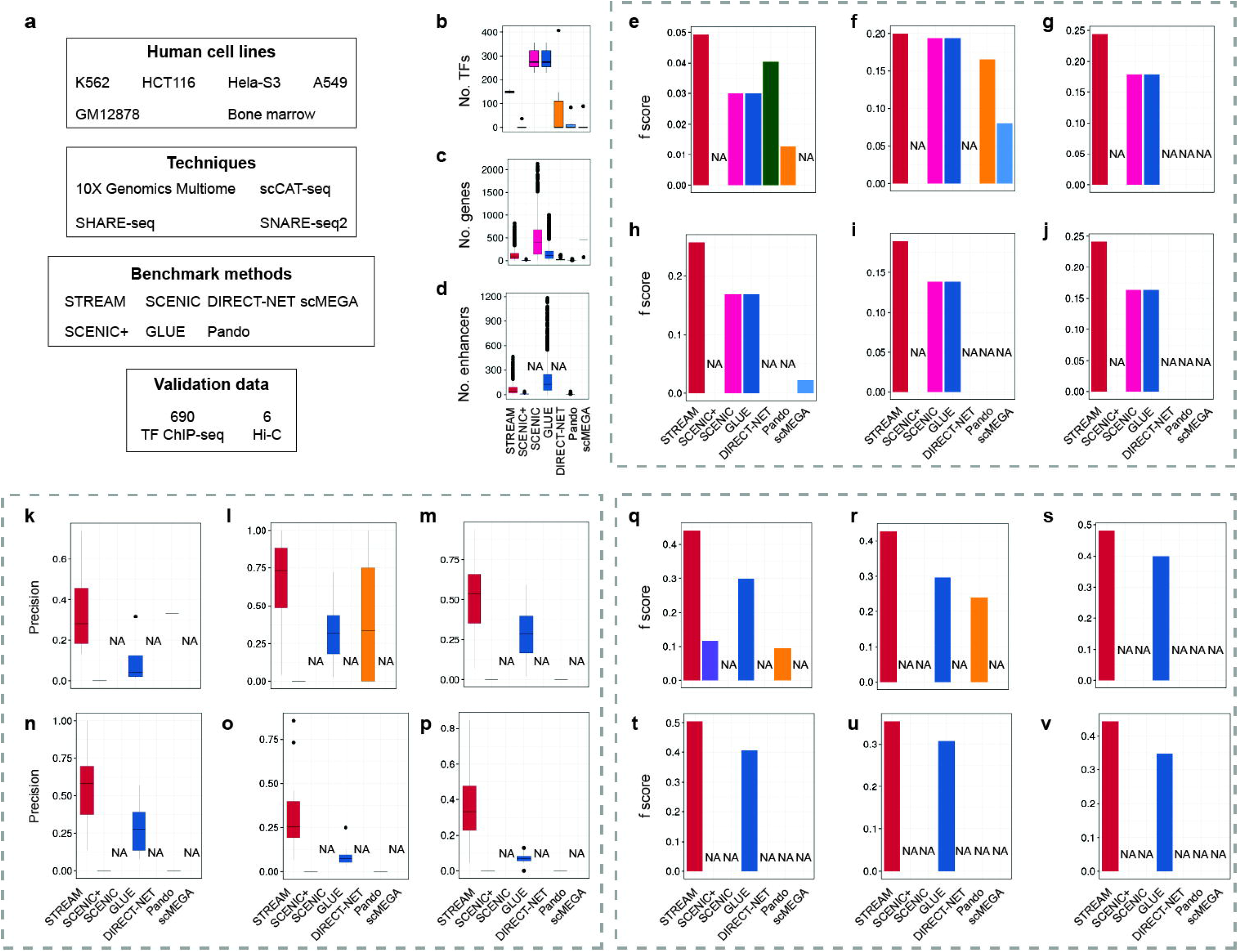
Evaluation of STREAM in comparison to other single-cell RNA-seq and ATAC-seq GRN inference methods using mainstream jointly profiled data. **a**. Chart of benchmarking strategy. **b-d**. Number of TFs (**b**) identified per method and distributions of the number of target genes (**c**), and enhancers per regulon and method (**d**). **e-j**. *f* score distributions from the comparison of TF recovery per method for datasets of human bone marrow (10X Multiome, **e**); mixture of K562, HCT116, and Hela-S3 (scCAT-seq, **f**); GM12878 (SHARE-seq, **g-h**); A549 (SNARE-seq2, **i**); and GM12878 (SNARE-seq2, **j**). **k-p**. Overlap between Hi-C links and predicted enhancer-gene relations per method for datasets of human bone marrow (10X Multiome, **k**); mixture of K562, HCT116, and Hela-S3 (scCAT-seq, **l**); GM12878 (SHARE-seq, **m-n**); A549 (SNARE-seq2, **o**); and GM12878 (SNARE-seq2, **p**). **q-v**. *f* score distributions from the comparison of regulon target genes, per method for datasets of human bone marrow (10X Multiome, **k**); mixture of K562, HCT116, and Hela-S3 (scCAT-seq, **l**); GM12878 (SHARE-seq, **m-n**); A549 (SNARE-seq2, **o**); and GM12878 (SNARE-seq2, **p**).

We evaluated STREAM’s performance against six specialized eGRN/GRN inference tools: SCENIC+^18^, SCENIC^2^, GLUE^6^, DIRECT-NET^3^, Pando^22^, and scMEGA^4^ (see **Methods** for details). Our findings showed that STREAM, SCENIC, and GLUE identified a range of 143-159, 229-356, and 229-356 TFs across the six datasets, respectively (**Fig. 3b**). In contrast, SCENIC+, DIRECT-NET, Pando, and scMEGA predicted 36, 89, 147-407, and 15-84 TFs across different datasets (**Fig. 3b** and **Supplementary Table 1**). A limitation of SCENIC+ was its inability to detect differentially accessible regions (DARs) in several datasets, preventing it from inferring eRegulons.

STREAM notably identified between 143 and 159 TFs, accounting for 70.4-78.3% of those with confirmed binding sites in JASPAR (203)^28^. In comparison, SCENIC+^18^, SCENIC^2^, and GLUE^6^, which utilize pycisTarget for TF-gene relation predictions, detected 36, 229-356, and 229-356 TFs respectively. Meanwhile, DIRECT-NET pinpointed 89 TFs^3^, Pando found between 147 and 407, and scMEGA recognized 15-84 TFs^4^. While GLUE determines enhancer-gene relationships through graph-linked unified embedding, it relies on SCENIC for inferring TF-gene associations^6^. This resulted in them predicting identical numbers of TFs, typically surpassing STREAM’s counts. On average, STREAM recognized 124 genes for each (e)regulon. In contrast, SCENIC+, DIRECT-NET, and Pando estimated 9, 28, and 2 genes per (e)regulon, respectively. Meanwhile, SCENIC, GLUE, and scMEGA identified 486, 157, and 403 genes per (e)regulon, respectively (**Fig. 3c**). Additionally, STREAM detected an average of 69 enhancers for each eRegulon. In contrast, SCENIC+, GLUE, and Pando estimated 10, 184, and 3 enhancers per eRegulon, respectively (**Fig. 3d**).

We subsequently assessed the biological relevance of the identified TFs by examining their alignment with those curated in 690 ENCODE TF ChIP-seq datasets specific to the same cell lines (**Fig. 3e-3j** and **Supplementary Table 5**). Our measure of relevance was the recovery rate of TFs involved in regulating (e)regulons. STREAM led the pack in terms of recovery, as demonstrated by its *f* score across the six datasets. DIRECT-NET followed suit, particularly shining in the 10X Genomics Multiome datasets, while SCENIC and GLUE were closely matched across other techniques. SCENIC+ managed to detect a limited set of eRegulons (36) in the 10X Genomics Multiome dataset (**Fig. 3e**), but strikingly, none coincided with the TFs curated from the ChIP-seq data of the corresponding cell line. While DIRECT-NET showed promise on the 10X Genomics Multiome (**Fig. 3e**), outperforming SCENIC and GLUE, its prowess waned on other platforms (**Fig. 3e-3j**). Turning to Pando, it exhibited a commendable performance on scCAT-seq, outperforming its counterparts (**Fig. 3f**).

Next, we gauged the accuracy of each method’s predicted target enhancers for every TF, referencing the TF ChIP-seq data from ENCODE (**Fig. 3k-3p**). For this comparison, we excluded SCENIC, DIRECT-NET, and scMEGA due to their inability to infer TF-enhancer relationships. Overall, STREAM’s predictions stood out for their precision, closely followed by GLUE (**Fig. 3k-3p**). Given that SCENIC+ failed to detect any TFs curated in the ENCODE TF ChIP-seq, its predictions for TF-enhancer relations did not correspond with ENCODE’s TF binding peaks (**Fig. 3k**). Mirroring the earlier TF recovery evaluation, Pando performs better than its peers on the scCAT-seq dataset (**Fig. 3l**).

For our final evaluation metric, we examined the accuracy of the predicted enhancer-gene relationships by cross-referencing them with chromatin interactions from deeply sequenced Hi-C data of the same cell lines (**Supplementary Table 6**). We excluded SCENIC, DIRECT-NET, and Pando for this assessment. Broadly speaking, STREAM consistently topped the charts with f scores ranging from 0.4 to 0.5 across all six datasets (**Figs. 3q-3v**), with GLUE closely following at scores between 0.3 and 0.4. SCENIC+ notched an *f* score of 0.12 on the 10X Genomics Multiome dataset, slightly outperforming Pando (**Fig. 3q**). The modest performance of SCENIC+ might be attributed to its limited detection of enhancer-gene relationships. Echoing the trends seen in our earlier evaluations of TF recovery and TF-enhancer relationship predictions, Pando stood out on the scCAT-seq dataset in comparison to the others (**Fig. 3r**).

In conclusion, STREAM consistently outperforms other methods in eRegulon inference across diverse cell lines and sequencing techniques. This performance is gauged by its effectiveness in TF recovery, accuracy in predicting TF-enhancer relationships, and proficiency in identifying enhancer-gene connections. The notable outcomes from STREAM arise from its use of the SFP model combined with submodular optimization. This approach provides a holistic representation of the connections between TFs and enhancers, as well as between enhancers and genes, rather than treating them in isolation.

### STREAM uncovers pseudotime-associated eRegulons and changing enhancer-gene relations alongside trajectories in AD

To showcase the capabilities of temporal scRNA-seq and scATAC-seq analysis, we utilize a case study by applying STREAM to an existing scRNA-seq and scATAC-seq dataset (with cell number *n* = 21,374). This dataset, composed of 32,286 genes and 66,861 enhancers, represents an AD mouse model brain studied over a time course (2.5 months: 7,517; 5.7 months: 2,357; and 13+ months: 11,500) and was generated using 10X Genomics Multiome. We discerned 27 cell clusters utilizing Seurat v.4.0.5, followed by manual annotation of each cluster through visualization of the expression and activity levels of marker genes (**Supplementary Table 2**). This process yielded seven distinct cell types: oligodendrocytes (Oligo), oligodendrocyte progenitors (OPC), inhibitory neurons (IN), excitatory neurons (EX), astrocytes (AG), microglia (MG), and endothelial & pericytes (**Figs. 4a-4b**). Upon deploying STREAM on this dataset, we identified 81 eRegulons of TFs with known relevance to AD, such as androgen receptor (AR)^29^, JUN family proteins^30^, ESR2^31,32^, FOSL2^33^, PLAG1^34^, RUNX1^35^, RORA^36^, and STAT2^37^, among others. We evaluated the similarity between eRegulons identified at different stages by calculating the fraction of overlapped enhancer-gene relationships, illustrated in **Fig. 4c**. A marked similarity was observed between pairs of eRegulons regulated by the same TF at varying stages. Additionally, some eRegulons, despite being regulated by different TFs, showed substantial intersection in enhancer-gene relationships. Building on these eRegulons, we identified cell-type-specific ones in EX, IN, and Oligo with eRegulon numbers of 18, 11, and 17, respectively.

**Fig. 4.**
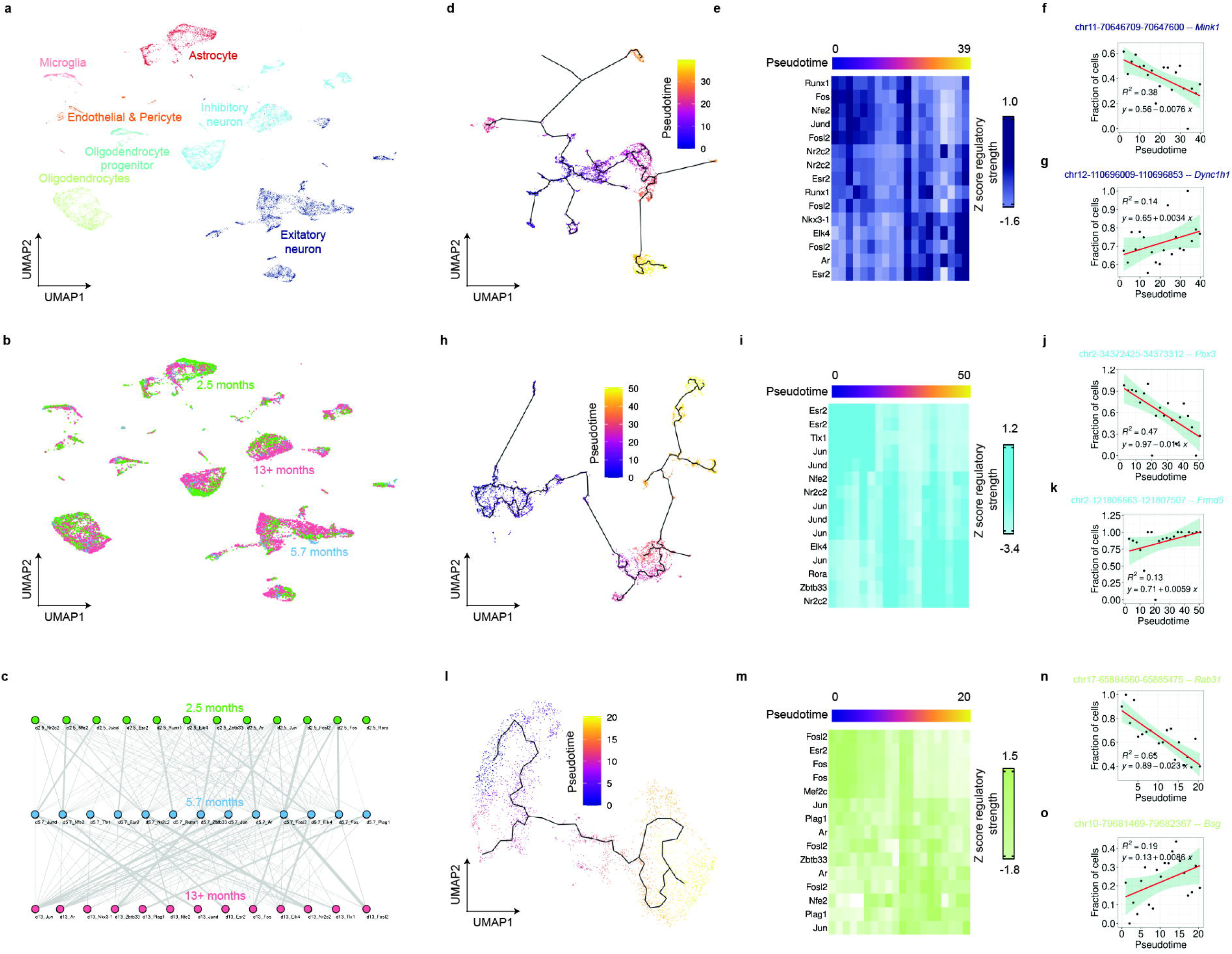
Analysis of Pseudotime-Associated eRegulons and Changing Trends of Enhancer-Gene Relations. **a-b**. UMAP plots color-coded by cell types **(a)** and stages **(b). c**. Graph visualizing eRegulon similarity identified at three stages, with nodes representing eRegulons and weighted edges indicating pairwise similarity defined as the Jaccard index of enhancer-gene relations of two eRegulons. **d**. UMAP plot with pseudotime color-coding in EX cells. **e**. Heatmap showing mean regulatory strengths of enhancer-gene relations in eRegulons specific to EX cells over pseudotime. **f-g**. Exemplary enhancer-gene relations demonstrating a monotonic trend in regulatory strengths over pseudotime in EX cells. Similar plots for IN cells **h-k** and Oligo **l-o**.

To examine the temporal dynamics of eRegulon regulatory strengths in EX, we isolated EX cells, inferred their developmental trajectory, and computed a pseudotime variable (**Fig. 4d**). By calculating the mean regulatory strength of enhancer-gene relationships within each eRegulon and converting these to *z*-scores, we observed notable temporal patterns (**Fig. 4e**). Herein, we define regulatory strength of an enhancer-gene relation as the proportion of cells in which the enhancer is accessible, and meanwhile, the gene is expressed among all cells belonging to a pseudotime/time period. Certain eRegulons showed monotonically changing regulatory strengths over pseudotime, with both increasing and decreasing trends evident. eRegulons regulated by RUNX1, FOS, NFE2, JUND, and FOSL2 exhibited a diminishing trend in gene expression, indicative of high regulatory strengths in cells at the start of the pseudotime trajectory. Meanwhile, elevated regulatory strengths were noted in cells positioned midway through pseudotime for eRegulons regulated by NR2C2, ESR2, RUNX1, and FOSL2. Furthermore, an uptick in regulatory strengths over pseudotime was detected in eRegulons governed by NKX3-1, ELK4, FOSL2, AR, and ESR2. These findings resonate with the known roles of these TFs in neurological contexts. For example, RUNX1 is known to exert a pro-neurogenic function in neural precursor cells^38^.

Diminished AR concentrations have been linked to cognitive deficits in aging and AD populations via pathway mediation^39^. Both JUN and FOS can influence apoptotic initiation and execution, implicating them in neuronal risk and cell death in AD^40,41^. Variations in ESR2 may correlate with increased AD susceptibility^42^. JUND, a member of the JUN protein family, can potentially impact cell apoptosis^40,41^. Additionally, ELK4 is associated with glia or neuron functions in AD 33, and FOSL2, part of the FOS protein family, takes part in gene regulation by forming the TF complex AP-1 with the JUN protein family^43^. We spotlighted the enhancer, chr11-70646709-70647600, bound by FOS and FOLS2, and noted its declining regulatory strength on Mink1 (**Fig. 4f**), a gene associated with cognition and critical in AD^44^. Furthermore, chr12-110696009-110696853, regulated by ELK4 and ESR2, displayed an increasing regulatory impact on the AD-associated gene Dync1h1 (**Fig. 4g**)^45^.

We extended our analysis to track changes in eRegulon gene expression over pseudotime in IN, using the same methodology applied to the EX cells (**Figs. 4h-4i**). Notably, we identified monotonic trends in the regulatory strengths of eRegulons within IN cells, which were controlled by ESR2, TLX1, JUN, JUND, NFE2, NR2C2, ELK4, RORA, and KAISO (encoded by Zbtb33).

Among these TFs, RORA is implicated in AD pathology and etiology^36^ and exhibits markedly upregulated expression in the AD hippocampus^36^. Additionally, KAISO is known to regulate central nervous system development^46^. Interestingly, ESR2, ESR2, and JUN were observed binding to chr2-34372425-34373312, regulating Pbx3, a significant gene in the central nervous system^47^. The regulatory strength of this interaction exhibited a decreasing trend over pseudotime (**Fig. 4j**). In contrast, we noted an ascending regulatory influence on Frmd5 from chr2-121806663-121807507, which was co-regulated by ELK4 and JUN (**Fig. 4k**).

Beyond neurons, we applied the same methodology to Oligo cells (**Fig. 4l**). We highlighted select eRegulons demonstrating monotonic regulatory strength changes over pseudotime (**Fig. 4m**).

These eRegulons were regulated by a roster of TFs similar to those identified in the EX (**Fig. 4e**) and IN (**Fig. 4i**) cell analyses. We honed in on two enhancers to further explore their regulatory influence on target genes. For instance, chr17-65884560-65885475, bound by FOSL2 and FOS, exhibited a declining regulatory impact on Rab31, which is often a target of RUNX1 in AD (**Fig. 4n**)^48^. Conversely, chr10-79681469-79682367 displayed an increasing regulatory strength on Bsg over pseudotime, under the governance of AR and JUN (**Fig. 4o**). Bsg is known to play a role in learning and memory^49^, and its dysregulation may result in sensory and memory function abnormalities^50^.

Broadly, by deducing the regulatory strength of eRegulons or enhancer-gene relationships across the developmental trajectory in EX, IN, Oligo, MG, AG, and OPC (**Fig. 4d-o and Supplementary Fig. 2, 3, and 4**), STREAM demonstrates its capacity to infer the critical role of eRegulons or enhancers in cellular developmental processes.

### STREAM unveils underlying eRegulons in diseased B cells in DSLL

To showcase the applicability of STREAM in cancer studies, we employed a DSLL dataset sequenced via 10X Genomics Multiome, encompassing 14,104 cells, 36,601 genes, and 70,469 enhancers. After conducting unsupervised clustering using Seurat v.4.0.5, we manually annotated cell types using an array of pre-established gene markers^51^. This dataset was subsequently finetuned to only encompass cells annotated by 11 cell types (**Fig. 5a**)^51^. Applying STREAM to the DSLL’s scRNA-seq and scATAC-seq dataset, we identified 50 eRegulons, comprising 19-290 genes and 9-70 active enhancers in 34-703 cells. Among these eRegulons, 99.6% of the genes (4,632) were associated with 1-10 enhancers, while 47.0% of the enhancers (4,658) were solely linked to the nearest gene (**Fig. 5b**).

**Fig. 5.**
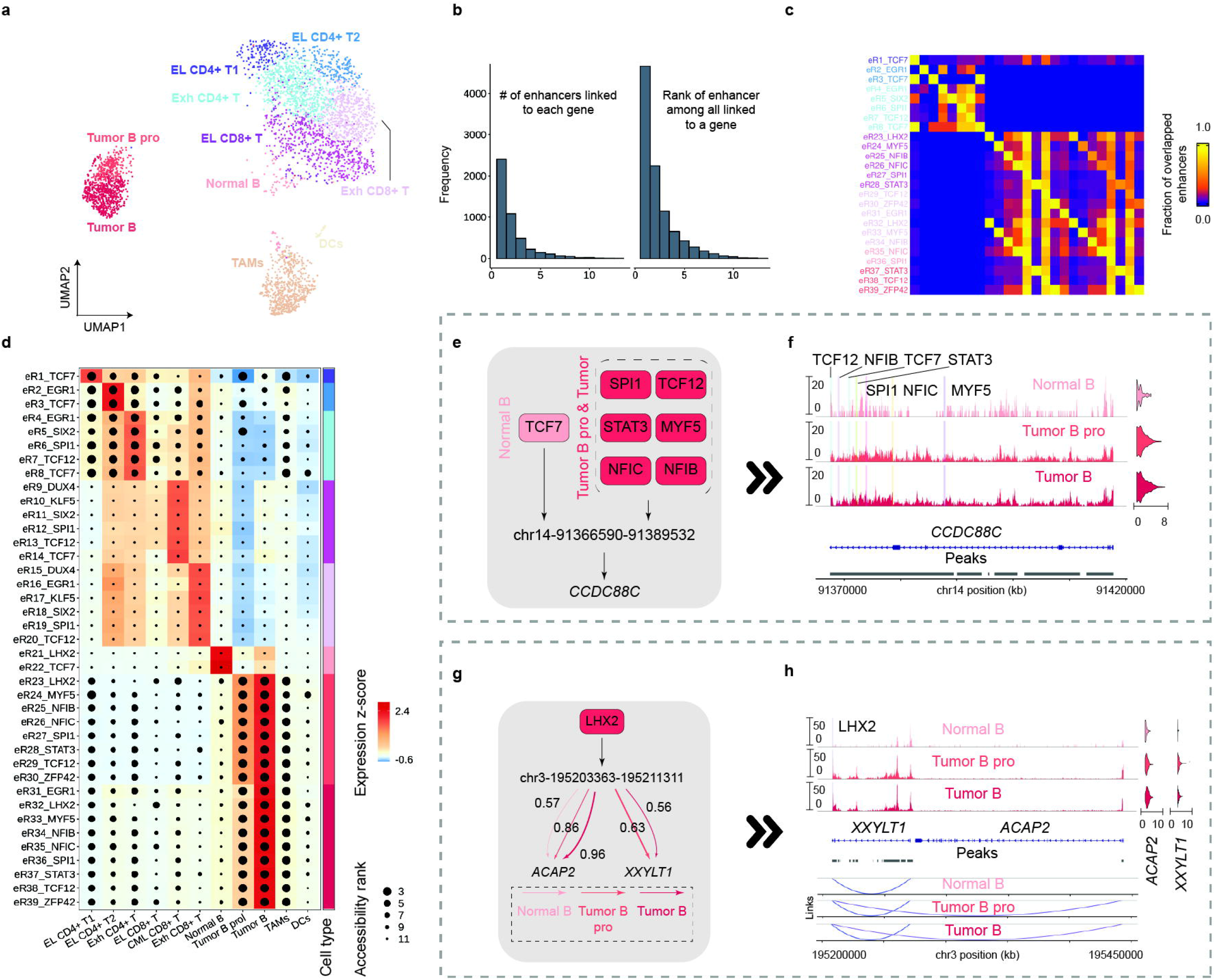
Key Enhancer and Enhancer-Gene Relations in B Cell Tumor Development Revealed by STREAM eRegulons. **a**. UMAP plot of cell types in DSLL. **b**. Distribution of the number of enhancers linked to each gene and rank distribution based oan n absolute distance between enhancer and gene. **c**. Overlap fraction of enhancers between eRegulon pairs, normalized by enhancer count in each row’s eRegulon. **d**. Heatmap-dotplot displaying eRegulon gene expression (color scale) and ranks of chromatin accessibility (size scale). **e**. Schematic showcasing TF variation regulating CCDC88C via chr14-91366590-91389532 binding among Normal B, Tumor B pro, and Tumor B cells. **f**. Chromatin accessibility profiles across three B cell types within chr14-91370000-91420000, labeled by TF binding sites from **e. g**. Schematic highlighting enhancer-gene relation variation between chr3-195203363-195211311 and gene pair (ACAP2 and XXYLT1) across three B cell types. **h**. Chromatin accessibility profiles within chr3-195203363-195211311 across three B cell types, labeled by LHX2 binding sites. Arcs show enhancer-gene links with color denoting the fraction of cells showing concurrent enhancer accessibility and gene expression.

Among the identified eRegulons, we isolated 37 that were specific to certain cell types, including Effector-like CD4+ T cell types 1 and 2 (EL CD4+ T1 and T2), Exhaustive CD4+ T cells (Exh CD4+ T), Effector-like CD8+ T cells (EL CD8+ T), Central memory-like CD8+ T cells (CML CD8+ T), Exhaustive CD8+ T cells (Exh CD8+ T), Dendritic cells (DCs), Normal B cells, Tumor B proliferation cells (Tumor B pro), and Tumor B cells. These eRegulons were exclusive to eight distinct cell types. A clear pattern of co-binding among TFs within these cell-type-specific eRegulons was observed (**Fig. 5c**). Moreover, we identified several well-known TFs regulating these cell-type-specific eRegulons in the DSLL dataset, including TCF7 and LHX2 in Normal B cells^52,53^, as well as SPI1^54,55^, TCF12^56^, STAT3^57,58^, NFIC^59^, NFIB^60^, and MYF5^61^ in both Tumor B pro and Tumor B cells. In many instances, we observed clear concordance between expression and chromatin accessibility changes in eRegulons governed by these TFs (**Fig. 5d**). However, in some cases, such as with eRegulons regulated by LHX2 (eR21), SPI1 (eR12), TCF7 (eR22), and TCF12 (eR13), we noticed that chromatin accessibility remained unchanged despite variations in expression. This underscores that chromatin accessibility can influence transcription within specific eRegulons across different cell types^62^.

To probe the underlying TF-enhancer-gene relationships in DSLL, we focused on cell-type-specific eRegulons within three cell types: Normal B, Tumor B pro, and Tumor B (**Figs. 5e-5h**).

We identified two, eight, and nine cell-type-specific eRegulons within Normal B, Tumor B pro, and Tumor B cells, respectively. In Normal B cells, we found two cell-type-specific eRegulons regulated by TCF7 and LHX2, respectively. These encompassed 14 and 73 genes regulated by TFs binding 11 and 82 enhancers, respectively. LHX2, coding for a protein that regulates the differentiation of multipotent lymphoid progenitor cells, is expressed early in B cell differentiation in a subepithelial lymphoid compartment^52^. TCF7, a vital TF for B cell development and maturation, well known for its role in early B cell lineage commitment and the progression of precursor cells within the bone marrow. Specifically, In our analysis, the importance of TCF7 in these cellular contexts was reaffirmed^52^. Additionally, in our examination of the Tumor B pro cells, we identified eight cell-type-specific eRegulons, which further underscores the molecular complexity of B cell differentiation and functionality. These eRegulons were regulated by a suite of TFs (LHX2, MYF5, NFIB, NFIC, SPI1, STAT3, TCF12, and ZFP42), which controlled 14-478 genes by binding 11-1,097 enhancers. For instance, MYF5, a member of myogenic regulatory factors, is key in myoblast specificity and cell proliferation^61^. Abnormal myoblasts accumulating in the bone marrow and crowding out healthy blood cells is a characteristic of leukemia/lymphoma^63^. NFIB and NFIC have been implicated in facilitating lymphoma cell proliferation, migration, invasion, and specificity^59,60^. TFs encoded by SPI1 and TCF12 were demonstrated to play roles in B cell differentiation and function^54-56^. Furthermore, ZFP42, encoding for REX1, is associated with pathways involved in lymph node oncogenesis^64^. The Tumor B cells showed a similar pattern to the Tumor B pro cells, with the additional presence of an EGR1 (Early growth response gene) eRegulon. EGR1, a known downstream effector of B-cell receptor signaling and Janus kinase 1 signaling in B-cell lymphoma, was discovered in Tumor B cells^65^. In summary, STREAM has demonstrated its capability in reliably uncovering TF-enhancer-gene relationships, thus providing insights into the biology of two Tumor B cell types.

In order to examine the influence of TF binding variation on critical gene expression across diverse B cell types, we focused on a marker gene of DSLL, CCDC88C, and a specific enhancer (chr14-91366590-91389532) through which CCDC88C was regulated by various TFs. Daple, encoded by CCDC88C, is known to modulate and activate Wnt signaling^66^, thereby reducing cell apoptosis^67^. Furthermore, CCDC88C has been found to be up-regulated in diseased B cells in lymphoma/leukemia compared to healthy ones^68^. We observed variation in TFs binding chr14-91366590-91389532 among three B cell types (**Fig. 5e**). Specifically, this enhancer was bound by TCF7 in Normal B cells, while it was co-bound by six different TFs (SPI1, TCF12, STAT3, MYF5, NFIC, and NFIB) in both Tumor B pro and Tumor B cells. Notably, we observed cooperative interactions among these TFs at this enhancer site (**Fig. 5f**). Finally, as illustrated in the violin plots, an up-regulation of CCDC88C between Tumor B pro or Tumor B cells versus Normal B cells was evident (**Fig. 5f**). In conclusion, chr14-91366590-91389532 can participate in the regulation of CCDC88C through different combinations of TFs, such as TCF7, SPI1, TCF12, STAT3, MYF5, NFIC, and NFIB. This leads to variations in CCDC88C expression and contributes to cell differentiation.

In addition to TF binding, we probed the influence of enhancer-gene relationships on marker gene expression. We selected two marker genes, ACAP2 and XXYLT1, both regulated by LHX2 through binding at chr3-195203363-195211311 (**Fig. 5g**). ACAP2 is a marker gene overrepresented in lymphoma^69^, while XXYLT1 is known to negatively regulate Notch signaling activation^70^, a pathway that can toggle an oncogene to a tumor suppressor^71^. We quantified the fraction of cells wherein the enhancer was accessible, and the gene was expressed, interpreting this as the regulatory strength. We found LHX2’s regulation of XXYLT1, through binding at chr3-195203363-195211311, was only evident in Tumor B pro and Tumor B cells with regulatory strengths of 0.63 and 0.56, respectively, but not in Normal B cells. Conversely, ACAP2 regulation by LHX2 through the enhancer at chr3-195203363-195211311 presented different regulatory strengths in the three cell types, i.e., Normal B: 0.57; Tumor B pro: 0.86; Tumor B: 0.96. A coverage plot incorporating chr3-195203363-195211311 and the two markers highlighted increased Tn5 insertion events in Tumor B pro and Tumor B cells compared to Normal B cells. This could suggest that the observed over-representation of marker expression, and consequent tumor development in B cells, may be driven by increasing regulatory strengths between chr3-195203363-195211311 and the two markers (**Fig. 5h**). This variation in regulatory strengths could be attributed to changes in chromatin accessibility. In conclusion, chr3-195203363-195211311 could be involved in regulating ACAP2 and XXYLT1 by LHX2 with varying regulatory intensities in different B cell types, resulting in changes in marker gene expression, cell functions, and differentiation.

### STREAM constructs comprehensive eGRNs showcasing the co-operativity among TFs in gene regulation across cell types

To present a comprehensive illustration of the various TF-enhancer-gene interactions in B cells within DSLL, we selected SPI1, STAT3, and TCF12 for further study of cooperative regulation and functional genomics analyses. ChIP-seq signal files for these TFs, obtained from the GM12878 cell line, were retrieved from the ENCODE project^72^. These selections allowed us to reconstruct an eGRN in Normal B, Tumor B pro, and Tumor B cells, respectively. SPI1 and TCF12 are critical for B cell development and commitment, respectively^54-56^, while STAT3 plays an active role in B cell lymphoma^57,58^. In addition, SPI1 and STAT3 have been found to work in concert to regulate several pathways crucial to tumorigenesis^73^. Similarly, TCF12, a member of the basic helix-loop-helix family, is implicated in promoting metastasis in various sites, including lymph nodes^74^. Moreover, these three TFs are involved in regulating CCDC88C, an activator of the Wnt signaling pathway. We highlighted their binding sites within the chr14-91370000-91420000 region across the three B cell types, as well as ChIP-seq datasets of SPI1, STAT3, and TCF12 in the GM12878 cell line (**Fig. 6a**). We observed significant overlaps between TF binding sites on the enhancer and ChIP-seq signals, particularly increased chromatin accessibility and ChIP-seq signals at regions commonly targeted by SPI1 and TCF12 (**Fig. 6a**). In conclusion, eGRNs inferred by STREAM can be effectively employed to examine the co-operativity among multiple TFs in gene regulation.

**Fig. 6.**
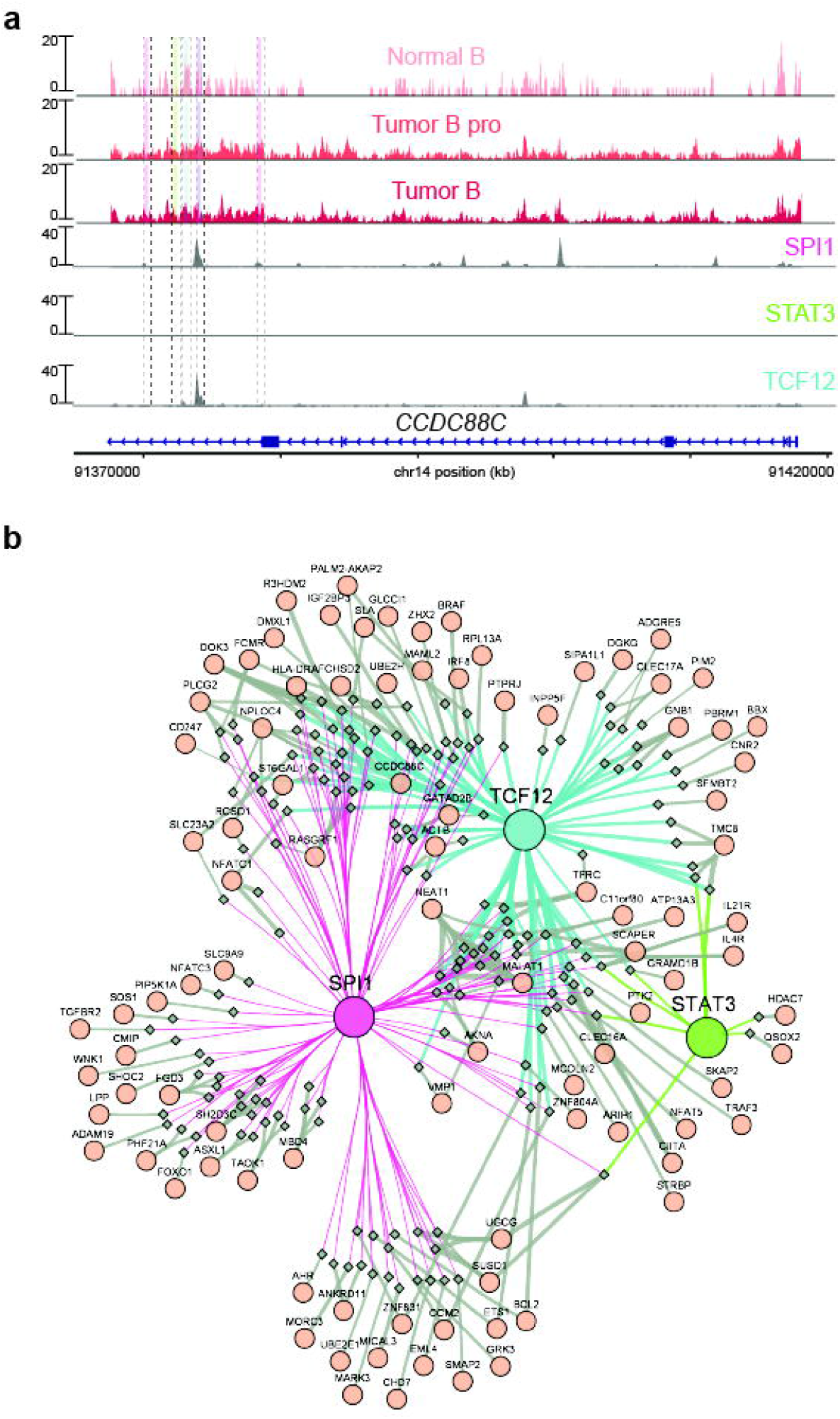
Interaction of SPI1, STAT3, and TCF12 in B cells. **a**. Demonstrates the chromatin accessibility profiles across three distinct B cell types, complemented by ChIP-seq signals for SPI1, STAT3, and TCF12 in the specified chromosomal region (chr14-91370000-91420000). **b**. Presents a visualization of an eGRN exclusive to proliferating Tumor B cells, constructed by the interplay of SPI1, STAT3, and TCF12 and confined to highly variable genes or enhancers. The thickness of the line connecting two nodes signifies the proportion of cells in which both nodes are either accessible (in the case of enhancers) or expressed (in the case of genes encoding TFs or their targets).

To provide a holistic depiction of the TF-enhancer-gene relationships in B cells within DSLL, we collated all TF-enhancer-gene relations centered on the three TFs of interest and their variable enhancers and genes. We then computed the weight of the TF-enhancer and enhancer-gene relations based on the fraction of cells in which the gene (target genes or gene encoding TFs) was active and its enhancer was accessible. This resulted in the creation of three distinct eGRNs, each composed of 276 nodes and 457 edges with varying edge weights across the three B cell types (**Fig. 6b** and **Supplementary Figs. 5-6**). In the eGRN specific to Tumor B pro cells, we identified 253 enhancers co-bound by SPI1 and STAT3 regulating 148 genes, 297 enhancers co-bound by SPI1 and TCF12 regulating 94 genes, and 208 enhancers co-bound by STAT3 and TCF12 regulating 69 genes, respectively (**Supplementary Table 3**). Pathway enrichment analysis against the KEGG database^75^ was performed for the three groups of regulated genes, yielding 45, 43, and 19 enriched pathways (*p*-value < 0.05 adjusted via Bonferroni correction), respectively (**Supplementary Table 4**). Genes co-regulated by these pathways exhibited higher regulatory strengths in Tumor B pro cells compared to Normal B and Tumor B cells. For example, genes co-regulated by SPI1 and STAT3 were enriched in several pathways, including Leukocyte transendothelial migration, Toxoplasmosis, Human papillomavirus infection, and Thyroid hormone signaling pathway. Each of these pathways has been implicated in lymphoma pathogenesis through various mechanisms such as immune dysregulation, the complication of lymphoma, increased risk due to persistent infection, and cancer cell proliferation stimulation^76-80^. Likewise, genes co-regulated by SPI1 and TCF12 were enriched in pathways such as Acute myeloid leukemia, Human cytomegalovirus infection, Phospholipase D signaling pathway, and Choline metabolism in cancer. These pathways have been associated with blood cancer development or progression, potential association with lymphoma, contribution to cancer progression, and lymphoma-associated dysregulation^80-85^. Finally, the pathways enriched by genes co-regulated by STAT3 and TCF12 were encompassed by those co-regulated by SPI1 and STAT3 or SPI1 and TCF12. Notable among these are Th1 and Th2 cell differentiation and Inflammatory bowel disease, both of which have been implicated in lymphoid cell development or differentiation and lymphoma risk, respectively^86^. In summary, the eGRNs inferred by STREAM can enable the exploration of the co-operativity among multiple TFs in gene regulation and their associated biological pathways.

## DISCUSSION

We introduce STREAM, an innovative framework designed to infer eGRNs from jointly profiled scRNA-seq and scATAC-seq data. Through rigorous benchmarking with six prominent (e)GRN inference methodologies across six distinct datasets, STREAM demonstrated superior performance in terms of TF recovery, TF-enhancer relation prediction, and enhancer-gene relation discovery. Moreover, the applicative power of STREAM has been substantiated through studies of an AD dataset over a time course and a DSLL dataset. On the AD dataset, our results revealed that STREAM can identify eRegulons in which genes display monotonic trends across pseudotime. STREAM subsequently computed the critical enhancer-gene relations exhibiting a monotonic trend quantified by the fraction of cells where the enhancer was accessible, and the gene was expressed. These predictions currently await experimental confirmation. STREAM’s application on the DSLL scRNA-seq and scATAC-seq dataset allowed us to pinpoint eRegulons specific to both healthy and disease-associated cell types. This case study revealed STREAM’s potential to unearth variations in TF binding and enhancer accessibility, indicating tumorigenesis. Furthermore, STREAM reconstructed eGRNs based on eRegulons across various cell types, thereby exploring the cooperative dynamics among TFs in gene regulation. These predictions also await experimental validation. In conclusion, STREAM has been shown to reveal pseudotime-associated eRegulons, unveil variations in enhancer-gene relations throughout cell lineages, identify the eRegulons underpinning diseased cell types, and elucidate the cooperative interactions among TFs in gene regulation. This array of capabilities positions STREAM as a potent tool, complementary to state-of-the-art tools, in the dissection of complex gene regulation systems.

Despite the novel insights and superior benchmarking performance STREAM offers, it comes with certain inherent limitations. First, STREAM leverages TF binding sites cataloged in the JASPAR database, which may constrain its ability to detect eRegulons governed by newly identified TFs. For instance, STREAM’s performance is sensitive to the level of consistency in hybrid biclustering. Second, STREAM may fall short when applied to scRNA-seq and scATAC-seq data with low-quality chromatin accessibility, exemplified by technologies such as Paired-seq^87^, which employs single-ended technology to sequence chromatin accessibility. Last, STREAM’s functionality is dependent on available annotations, specifically TF binding sites from JASPAR^88^. Therefore, its applicability for constructing eGRNs is limited to the range of TFs included in the JASPAR database.

In summary, STREAM offers a powerful platform for eGRN inference, elucidating complex regulatory mechanisms from multiple modalities across a variety of application scenarios. The methodological framework underpinning STREAM can be further enhanced for the identification of eRegulons, eGRNs, and cell types, drawing upon strategies from cell-type-specific or differential GRN prediction approaches such as DeepMAPS^51^, GLUE^6^, scMTNI^89^, and sc-compReg^90^, etc. This integrative strategy could pave the way for deeper exploration and comprehension of cellular dynamics and GRNs.

## METHODS

### The STREAM framework

#### Step 1: FGM prediction

FGM is a highly structured expression pattern of a gene set, which tend to be functionally related or co-regulated in a particular cell subpopulation^91^. To identify FGMs, STREAM accepts as input an expression matrix (*X*_*n*×*o*_) and a chromatin accessibility matrix (*Y*_*m*×*o*_), both post-quality control (QC). These matrices represent the expression levels (numerical) of *n* genes and chromatin accessibility (binary) of *m* peaks, respectively, across *o* cells (**Fig. 2a** and **Supplementary Fig. 1a**). To extract the underlying gene regulatory information from gene expression, the expression matrix *X*_*n*×*o*_ is transformed into a discretized matrix 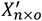 via a left truncated mixture Gaussian (LTMG) model^92^, known as the LTMG matrix,. This LTMG model capably captures the diversity of gene expression states regulated by varying transcriptional inputs across individual cells^92^. Following this, the sRNA-seq analysis tool, IRIS-FGM, is applied to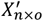]to obtain biclusters *B*_*k*_, where *k* = 1, …, *l*. Each bicluster, for example, *B*_*k*_, signifies an FGM^91^. These FGMs will act as the input to **Step 2**.

#### Step 2: SFP model

The STREAM methodology involves an SFP model, designed to reverse-engineer the underlying eGRNs by extracting combinations of enhancer-gene and enhancer-enhancer relations that are most likely to lead to FGMs (**Fig. 2a** and **Supplementary Fig. 1b**)^91^. For each FGM, *B*_*k*_, an LTMG submatrix 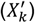 and a chromatin accessibility submatrix (*Y*_*k*_) are constructed. The rows of these matrices correspond to the genes of *B*_*k*_ and all the enhancers, respectively. The columns of both 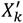 and *Y*_*k*_ are restricted to the cells present in *B*_*k*_. Utilizing Signac v.1.10.1^93^ and Cicero v.1.17.2^94^, we construct a heterogeneous graph, *G*^(*k*)^ = (*V*^(*k*)^, *E*^(*k*)^), where *V*^(*k*)^ represents the rows from 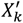 and *Y*_*k*_. The edges in this graph signify both enhancer-enhancer and enhancer-gene relationships based on 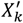 and *Y*_*k*_. This construction is used to compute the enhancer-gene edges and enhancer-enhancer edges, respectively. The “cost” of enhancer-gene edges is weighted by the absolute value of Pearson correlation coefficients, while enhancer-enhancer edges are weighted by the absolute value of covariances. This involves subtracting the minimum-maximum normalized edge weight from one. The set of nodes that denote genes in *V*^(*k*)^, are then divided into several subsets, 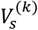 where *s* = 1, …, *t*. These subsets ensure that gene nodes belonging to the same subset are connected by a path in *G*^(*k*)^. These subsets are referred to as terminal sets. In the subsequent sections, the distinction between genes/enhancers and nodes representing them in graphs is not made.

In the context of the weighted undirected graph *G*^(*k*)^ and *t* terminal sets 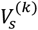, the aim of the SFP model is to identify a forest *F*^(*k*)^ with minimum cost within *G*^(*k*)^, also known as the Steiner Forest (SF). This forest ensures that the nodes within each 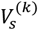 are connected via the edges of the forest. Given the NP-hard nature of the SFP problem, a heuristic approach is used for its resolution. It is clear that *F*^(*k*)^ consists of one or more trees 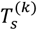, through each of which nodes within each terminal set are interconnected. *F*^(*k*)^ is identified by discovering each 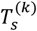 individually. For each terminal set 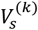, the process begins with the pair of genes that are co-regulated across the maximum number of cells. The shortest path in *G*^(*k*)^ between them is then determined. This shortest path serves as the initial 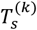. Next, the process involves: (*i*) selecting the gene from 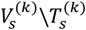 that results in the shortest path to 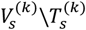 and (*ii*) adding nodes of the shortest path to 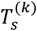. This continues until 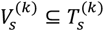 Finally, all 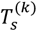, *s* = 1, …, *t* are merged to obtain the SF, denoted as *F*^(*k*)^,

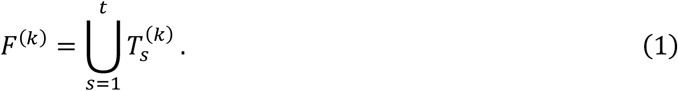

Within the structure of *F*^(*k*)^, we identify enhancer-gene relations and categorize them into subsets based on the TF binding sites as recorded in JASPAR^88^. In this way, all enhancer-gene relations within each subset are associated with the same TF. We then retain only those enhancer-gene relation subsets which are linked to a minimum of two genes.

Next, we transform each enhancer-gene subset into an initial HBC. This corresponds to a pair of biclusters, each representing a subset of genes 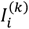, a subset of enhancers 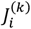, and a subset of cells 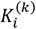, which we define as an HBC,

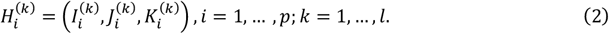

In each HBC 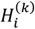, we sequence the cells in 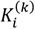in descending order based on their average gene expression values across 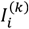, resulting in a monotonic trend 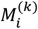. Furthermore, for each enhancer in 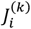, we calculate the average ratio 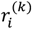of cells where the enhancer is accessible compared to all cells in 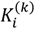. We define the minimum of 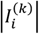and 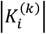as the HBC score, on which we consolidate and rank all HBCs in decreasing order. These ranked HBCs will serve as seed sets for hybrid biclustering in **Step 3**, denoted as

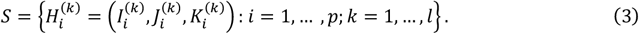

#### Step 3: Hybrid biclustering

Starting with the seeds predicted in **Step 2**, STREAM utilizes hybrid biclustering to simultaneously identify co-regulated genes and co-accessible enhancers in a specific cell subpopulation. Based on a seed in *S*, we expand it in both the vertical (genes and enhancers) and horizontal (cells) dimensions, ensuring the monotonic trend of gene expression across cells within the HBC is maintained (**Fig. 2b** and **Supplementary Fig. 1c**). Hybrid biclustering, which is equivalent to simultaneous biclustering on two matrices, is a computationally challenging problem due to its NP-hard nature^95^. Therefore, we utilize a heuristic approach to perform hybrid biclustering. If an HBC can no longer be expanded, it is output as the final HBC. The details are as follows. If *S* is empty, the hybrid biclustering process stops; otherwise, the first HBC in *S* is checked to see if it is a qualified seed. If it is not, it is removed from *S*, and this step will be repeated; otherwise, the seed is expanded by iteratively finding the longest path from directed acyclic graphs (DAGs) that represent the monotonicity of gene expression among cells. In this context, a seed is deemed qualified if it shares less than a default α (α = 0.50) proportion of genes with any previously identified HBCs.

For the current HBC 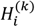we construct a DAG, 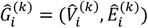where nodes represent cells in 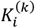and each directed edge connects a higher-ranking cell to a lower-ranking cell according to the monotonic trend 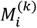To avoid infinite long paths in the DAG, pairs of nodes within a loop or circle are condensed into a single node. Hybrid biclustering aims to enhance the HBC score, denoted as

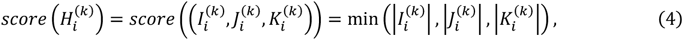

while upholding the monotonic trend 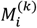among the cells of 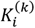we construct a candidate gene set in each iteration. This set includes genes that: (*i*) are not part of 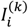and, (*ii*) are linked to at least one enhancer bound by the same TF as 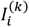, which we term as supporting enhancers for the gene. For each gene *g* within the candidate gene set, we create a DAG, 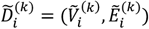 by: (*i*) excluding the cells from 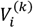where *g* is not expressed and concurrently eliminating their corresponding edges from 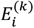 (*ii*) removing the edges from 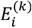that violate the monotonic trend of expression values of *g* among cells in 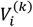and (*iii*) removing the genes which lack supporting enhancers accessible in 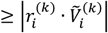 cells. Using dynamic programming, we identify the longest path from each 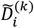 and single out the gene (optimal gene) that provides the longest path 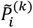 among all genes in the candidate gene set^96^. We continue to use 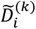 to denote the DAG crafted based on the optimal gene. Following this, we generate a new HBC,

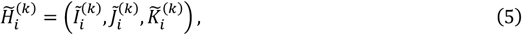

in the following manner: (*i*) we assign 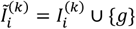 (*ii*) if no supporting enhancer of *g* is included in 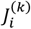we designate 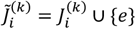 where *e* represents the supporting enhancer of *g* that is accessible in the maximum number of cells of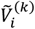 If there are supporting enhancers in 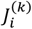, we keep 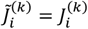Subsequently, (*iii*) we set 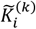as the nodes in 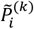. If the score of 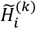 is greater than or equal to the score of 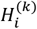we assign 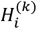as 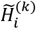 and 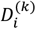 as 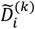Otherwise, we terminate the iteration, output 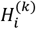and eliminate the current seed from *S*. If *S* is empty, we halt the hybrid biclustering. If not, we select the next qualified seed for hybrid biclustering.

In conclusion, we arrange the HBCs in the descending order based on the HBC score.

Furthermore, within each HBC, we establish enhancer-gene relations 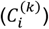by connecting each gene to the enhancers situated within a 250 kb distance from the transcription start site (TSS)^21^. Correspondingly, we denote each HBC as

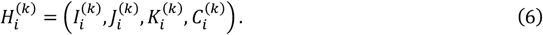

We send all the HBCs for submodular optimization in **Step 4**.

#### Step 4: HBC optimization

The optimization of HBCs aims to identify a subset of HBCs that yield not just promoted diversity but also reduced redundancy. This subset will be output as the set of eRegulons. The solution to HBC optimization comprises three key components (**Fig. 2c** and **Supplementary Fig. 1d**): (*i*) an evaluation function that establishes the number of eRegulons in a dataset; (*ii*) an objective function that quantifies the informativeness of a subset of HBCs; and (*iii*) a submodular optimization algorithm that discerns a subset of HBCs that ranks highly based on the objective function^24^.

In our model, the evaluation function is formulated as follows:

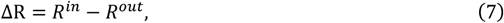

Here, *R*^*in*^ refers to the total number of enhancer-gene pairs connected through the relationships outlined in HBCs, whereas *R*^*out*^ signifies the number of enhancer-gene pairs located within a default proximity of 250 kb from each other, but lack any established connection through the enhancer-gene relationships present in the HBCs.

Adopting the facility location objective function, we assess the fraction of data available in the comprehensive HBC set that is encapsulated within the HBC subset. Let us denote *U* as the full HBC set and *W* as the subset of HBCs. Then, the facility location function *f*, mapping from the power set of *U* to the real numbers, is expressed as follows:

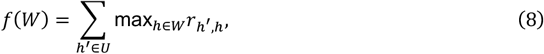

where *rh*′ *h* determines the pairwise similarity between HBCs *h*′ = (*I*′, *J*′;, *K*′, *C*′) and *h* = (*I, J, K, C*), and is given by:

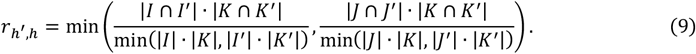

Intuitively, the facility location function achieves a high value when every HBC in *U* has at least one representative in *W* that is similar. We define the conditional gain of *f* as:

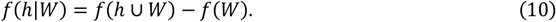

Submodular optimization initiates with the empty set *W*_0_ = ∅ and at each iteration *i*, it adds the HBC *h*_*i*_ that maximizes the conditional gain *f*(*h*_*i*_ |*W*_*i*-1_) with ties broken arbitrarily (i.e., identifying 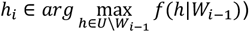then proceeds to update *W*_*i*_ ← *W*_*i*-1_ ∪ {*h*_*i*_}. The algorithm ceases when the number of HBCs to select is met. We optimize the number of HBCs by maximizing the difference between the number of enhancer-gene relations within HBCs and that between different HBCs. We subsequently output these HBCs as eRegulons.

### Benchmarks

#### Datasets

For benchmarking analyses, this study collated a total of six previously published datasets encompassing both scRNA-seq and scATAC-seq data from human specimens encompassing six cell lines (see **Supplementary Table 1**). These datasets were obtained using four different technologies: 10X Genomics Multiome, scCAT-seq^25^, SHARE-seq^26^, and SNARE-seq2^27^. These are publicly accessible via the NCBI Gene Expression Omnibus (GEO), the official website of 10X Genomics, or within the relevant published literature. For our case studies, we selected a 10X Genomics Multiome dataset derived from coronal sections of an AD mouse model brain hemisphere, sampled at 2.5, 5.7, and over 13 months. Furthermore, we included a 10X Genomics Multiome dataset comprising approximately 14,000 sorted nuclei sourced from lymph nodes in a DSLL model.

#### Preprocessing scRNA-seq and scATAC-seq datasets

For all the scRNA-seq and scATAC-seq data employed in this study, we loaded count matrices into the program using Read10X_h5, read_10x, and read.table functions, respectively. For scRNA-seq, the count matrix was converted into a Seurat object using the CreateSeuratObject function from Seurat (v.4.0.5). The percentage of mitochondrial RNAs was calculated using the PercentageFeatureSet function. For scATAC-seq, enhancers belonging to standard chromosomes were retained using the standardChromosomes function. We then generated genomic annotations using the GetGRangesFromEnsDb function, with the parameter ensdb set to EnsDb.Hsapiens.v86 (for hg38) and EnsDb.Hsapiens.v75 (for hg19), respectively. For all datasets, we used the intersect function to retain the common cells present in both the scRNA-seq and scATAC-seq matrices. For the 10X Genomics Multiome datasets, the fragments of scATAC-seq were incorporated into the Seurat objects via the CreateChromatinAssay function. Finally, we used the subset function in Seurat for QC, based on the percentage of mitochondrial RNA as well as the count of scRNA-seq and scATAC-seq.

#### Benchmark methods

We conducted a comprehensive comparison of STREAM with six methods for (e)GRN inference. The compared methods were as follows: (*i*) SCENIC (pySCENIC v.0.12.1), a rapid Python-based implementation of the SCENIC pipeline allows biologists to deduce TFs, GRNs, and cell types from scRNA-seq data with unparalleled speed^2^; (*ii*) SCENIC+ v.1.0.1, a Python package designed to construct eGRNs from either combined or individual datasets of scRNA-seq and scATAC-seq^18^; (*iii*) GLUE v.0.3.2, a modular deep learning framework for jointly analyzing scRNA-seq and scATAC-seq data and inferring regulatory interactions simultaneously^6^; (*iv*) DIRECT-NET v.1.0.0, an R toolkit for inferring CREs and constructing GRNs from parallel scRNA-seq and scATAC-seq data or only scATAC-seq data^3^; (*v*) Pando v.1.0.5, an R package that leverages multi-modal single-cell measurements to infer GRNs using a flexible linear model-based framework^16^; (*vi*) scMEGA v.1.0.1, an R package designed to infer GRNs based on Seurat, Signac, and ArchR^4^. To ensure a fair comparison, we followed the tutorials and recommendations provided by each method.

#### Workflow introduction of SCENIC

The SCENIC workflow is a comprehensive framework designed for inferring GRNs using scRNA-seq data. It integrates three packages, namely GENIE3^97^, RcisTarget (pycisTarget)^2^, and AUCell^2^, to collectively analyze and interpret the underlying regulatory interactions. pycisTarget is the Python version of RcisTarget which boasts the most expansive motif collection to date, encompassing 32,765 unique motifs gathered from 29 diverse collections. GENIE3 is employed in SCENIC to uncover potential TF-gene relations by identifying gene co-expression patterns. This approach leverages the assumption that genes with similar expression profiles will likely be regulated by the same set of TFs^97^. RcisTarget (pycisTarget), another crucial component of SCENIC, enables the identification of direct TF-gene relations by performing motif enrichment analysis. By comparing known TF binding motifs with the regulatory regions of target genes, RcisTarget predicts the potential TFs that regulate specific genes within the scRNA-seq data^2^. Lastly, AUCell is utilized in SCENIC to quantify the activity of regulons in single cells. Regulons are defined as sets of co-regulated genes controlled by the same TFs. AUCell assesses the activity of these regulons across individual cells, providing valuable insights into the regulatory dynamics within the scRNA-seq dataset^2^. By integrating these three packages, SCENIC offers a comprehensive approach to infer GRNs from scRNA-seq data, providing a valuable resource for the identification and characterization of regulatory interactions in single-cell transcriptomic studies.

#### Framework description of SCENIC+

SCENIC+ is an advanced computational workflow designed to construct eGRNs through a structured three-step process^18^. Initially, to pinpoint candidate enhancers, it processes scATAC-seq data utilizing pycisTopic, a Python reiteration of cisTopic^98^. This step employs DARs and topics, sets of co-accessible chromatin regions, to identify potential enhancer candidates across varying cell types or states, with topics proving more enriched for functional enhancer regions than DARs. The second phase focuses on unearthing potential TF binding sites nestled within these candidate enhancers. For this, SCENIC+ embarks on a motif enrichment journey, harnessing the power of pycisTarget. These motifs, spanning multiple species, are clustered based on similarity. Scoring all motifs within a cluster has shown to offer superior precision and recall compared to using singular archetype motifs. The analysis is further strengthened by employing two robust algorithms, cisTarget and the differential enrichment of motifs (DEM), outperforming the existing tool, Homer. DEM excels in discerning differential motifs within regions of analogous motif content. In the final phase, SCENIC+ leverages GRNBoost2 to ascertain the significance and regulatory direction (either activating or repressing) of both the TFs and enhancer candidates in relation to target genes. By seamlessly combining motif enrichment results with GRNBoost2 outputs and conducting a secondary enrichment analysis, the workflow precisely pinpoints the most fitting TF for each motif set. This culminates in the creation of the eRegulon, which is a composite of a TF along with its cohort of target enhancers and genes.

#### Model overview of GLUE

The GLUE framework is a modular deep learning-based approach designed to integrate scRNA-seq and scATAC-seq data for inferring *cis*-regulatory relations^6^. The workflow in GLUE begins with QC of the scRNA-seq and scATAC-seq data. The final Seurat objects, representing the processed and quality-controlled data, are then converted into the standard file format of 10X Genomics (Feature-Barcode Matrices). This conversion ensures compatibility with the GLUE framework. GLUE employs variational autoencoders (VAEs) to model cell states as low-dimensional cell embeddings for each modality^99^, namely scRNA-seq and scATAC-seq. These VAEs capture the underlying biological information and reduce the high-dimensional data into a more manageable representation. To integrate the two modalities, GLUE utilizes prior biological knowledge about the locations of enhancers and genes to construct a guidance graph. This graph provides a framework for aligning and integrating the enhancer-gene relations inferred from scATAC-seq and scRNA-seq data. The alignment process in GLUE involves iterative adversarial multimodal alignment, where the enhancer-gene relations are refined using the guidance graph. This iterative refinement helps to improve the accuracy of the inferred *cis*-regulatory relations. Finally, GLUE leverages the SCENIC (pySCENIC) method to construct TF-gene relations^2^. By filtering out the enhancers that are linked to the target genes of a TF, the TF-enhancer-gene relations can be obtained. Through the integration of scRNA-seq and scATAC-seq data, as well as the utilization of VAEs, guidance graphs, and iterative alignment, GLUE offers a comprehensive framework for inferring *cis*-regulatory relations and providing insights into the interplay between TFs, enhancers, and genes in cellular regulatory processes.

#### Pipeline introduction of DIRECT-NET

DIRECT-NET is a computational tool designed to identify functional CREs, connect them to their corresponding target genes, and subsequently assemble TF to CRE to construct GRNs^3^. The tool begins by classifying any peak within 500 bp upstream of a gene’s TSS as the promoter, while other open chromatin areas outside the promoter but within a set boundary (typically 250 kb on both sides) are designated as distal candidate functional regions. The input for DIRECT-NET can either be paired scRNA-seq and scATAC-seq data or just scATAC-seq data. Recognizing the sparsity of single-cell sequencing data, especially binary-like scATAC-seq data, the first step involves aggregating binary epigenomic signals or combined transcriptomic and epigenomic profiles across similar cells, inferred via nearest neighbors in a data-derived low-dimensional space. Subsequently, DIRECT-NET discerns CREs by regressing the gene expression level (if both data types are present) or the promoter’s accessibility score (if only scATAC-seq data), against the accessibility scores of potential peaks nearby using the XGBoost model. Functional CREs are then pinpointed through the importance scores derived from XGBoost. Ultimately, a TF-gene regulatory network is established by integrating predicted functional CREs with known motif patterns from public databases to deduce CRE-TF and CRE-gene associations.

#### Methodological details of Pando

Pando constructs a GRN by first identifying accessible regulatory regions, using conservation information and prior CRE annotations^16^. These regions are assigned to nearby genes, and TF binding sites are predicted within each region. Pando then employs a regression model to determine the relationship among the expression of target genes, TF expression, and TF binding site accessibility. As a result, it simultaneously identifies groups of genes either positively or negatively regulated (gene modules) and the associated regulatory genomic areas (regulatory modules) for each TF.

#### Framework introduction of scMEGA

scMEGA is designed for GRN inference through three main steps^4^: (1) multimodal data integration, (2) candidate TFs and gene identification, and (3) GRN construction. For jointly profiled scRNA-seq and scATAC-seq data, the first step can be skipped. Using integrated data, scMEGA identifies candidate TFs and genes by mapping a pseudotime trajectory with the ArchR package^21^. Chromatin accessibility profiles assist in estimating each TF’s binding activity, and the tool correlates this with gene expression to pinpoint active TFs. Along the pseudotime trajectory, scMEGA selects top variable genes and correlates them with peak accessibility. A quantitative GRN is formed based on TF binding activity and gene expression. Enhancers, defined as peaks at least 2kb from a gene’s TSS, are used to link TFs with genes. Combining the enhancer-based and quantitative GRNs, scMEGA produces a final eGRN, with interactions weighted by correlation strength.

### Evaluation metrics of eGRN inference

We evaluate the effectiveness of various eGRN/GRN identification techniques, and examined their performance following SCENIC+ using three perspectives: TF recovery, TF-enhancer relation prediction, and enhancer-gene relation discovery^18^.

#### TF recovery

To evaluate whether the methodologies in place effectively pinpointed the relevant TFs for the distinct cell lines, we sourced all the TF ChIP-seq bed files specific to these six cell lines from the ENCODE database (**Supplementary Table 5**). With these datasets in hand, we proceeded to analyze each method’s accuracy. For every dataset, we identified overlaps between the TFs that were present in the (e)GRNs and those listed in the ENCODE TF ChIP-seq data. Such overlapping instances were classified as true positives. After establishing these overlaps, our next step involved the computation of f scores. This metric was specifically chosen as it allowed us to gauge and compare the effectiveness of TF recovery across the different methodologies we were assessing. Through this detailed procedure, we aimed to establish a comprehensive understanding of each method’s capability to accurately identify and recover TFs for the given cell lines.

#### TF-enhancer relation prediction

To conduct an evaluation of the TF to enhancer associations deduced by the methods being benchmarked, we took the initial step of acquiring all available TF ChIP-seq bed files from the ENCODE database, specifically tailored for these six distinct cell lines (**Supplementary Table 6**). For each cell line and their corresponding TFs, it was essential to ensure that we used the most comprehensive data. Thus, out of the multiple bed files, we opted for the one encompassing the highest number of signal peaks related to that specific TF. Once this selection was made, our next task was to determine the degree of correspondence between the predicted TF-binding enhancers as per our methodologies and the TF-binding peaks already documented in the ENCODE ChIP-seq. Given the potentially vast number of ChIP-seq peaks present in the ENCODE database, we felt it was necessary to employ a precision metric. By leveraging precision, we aimed to effectively evaluate the accuracy and relevance of the predicted TF-enhancer associations, ensuring that our predictions were not only high in number but also of significant quality and relevance.

#### Enhancer-gene relation discovery

To evaluate the accuracy of the inferred enhancer-gene connections as deduced by the various methods under consideration, we utilized the Hi-C data corresponding to the six specific cell lines, sourced from the ENCODE database. Our first step involved sifting through the available chromatin contact matrix files for each cell line. Given the different sizes of these files, we ensured that we selected the matrix file with the largest dataset, stored in the .hic format. To make sense of this .hic format and to convert it into a more usable contact matrix, we employed the straw function available in strawr v.0.0.91. The resultant contact matrix displayed rows and columns as bins, each bin being 2,500 kb in size. The individual matrix values represented read counts, providing an indication of the frequency of contact between specific chromatin regions. In parallel, our goal was to pinpoint the exact locations of genes. For this purpose, we accessed gene annotations from several R packages. Specifically, we used EnsDb.Hsapiens.v86 for the hg38 genome version and EnsDb.Hsapiens.v75 for hg19. Once equipped with this data, our next task was to determine the extent of overlap between our predicted enhancer-gene associations and the contact data between bin-to-bin derived from the ENCODE Hi-C datasets. To quantify how accurately our methods predicted the enhancer-gene connections, we computed the *f* scores as a metric to assess our predictions.

### Prediction of enhancer-gene relations associated with the trajectory in AD

Cell-type-specific trajectories were inferred from the transcriptomics component of both the scRNA-seq and scATAC-seq data. This was accomplished by employing Monocle3 v.1.0.0, with cells labeled as being at the 2.5-month stage serving as the root of each trajectory^100^.

### Cell-type-specific eGRN construction in DSLL

We established a cell-type-specific eGRN for the DSLL cell type through a two-step process. First, we identified cell-type-specific eRegulons via a hypergeometric test (with a *p*-value threshold of < 0.05), subsequently adjusting for multiple hypothesis tests using the Bonferroni correction. If the cell set in which an eRegulon is active is significantly enriched in a cell type, we referred this eRegulon specific to the cell type. We constructed cell-type-specific eRegulons by merging eRegulons regulated by the same TF specific to the same cell type. In the second step, we assembled the cell-type-specific eGRN by amalgamating all the cell-type-specific eRegulons and linking all TF-enhancer-gene relationships. We used the FindMarkers function in Seurat v.4.0.5, under default parameters, to uncover differentially expressed genes (DEGs) or DARs that were overrepresented in each cell type. The DEGs/DARs with adjusted *p*-values < 0.05 were identified as significant.

## DATA AVAILABILITY

All datasets analyzed in this study were published previously. The corresponding descriptions and pre-processing steps are described in the **Methods**.

## SOFTWARE AVAILABILITY

Detailed tutorials and documentation on the STREAM workflow are available at https://github.com/YangLi-Bio/stream2. Scripts to reproduce the analyses presented in this manuscript are available at https://github.com/YangLi-Bio/stream2_analyses.

## ACKNOWLEDGEMENTS

This work was supported by award R01GM131399 from the National Institute of General Medical Sciences of the National Institutes of Health. The work was also supported by the award NSF1945971 from the National Science Foundation. This work was supported by the Pelotonia Institute of Immuno-Oncology (PIIO). The content is solely the responsibility of the authors and does not necessarily represent the official views of the PIIO. The authors thank Qiuqin Wu, Xiaoying Wang, Zhenyu Wu, Yujia Xiang, Shuangquan Zhang, and Cindy Tong for their assistance in data collection, pipeline development, and tool testing.

## AUTHORS’ CONTRIBUTIONS

Q.M. and B.L. conceived the basic idea and designed the overall analyses. H.F. provided a guide for biological annotation in the AD case study. Y.L. designed the specific experiments and developed the framework. Y.L., A.M., Y.W., Q.G., and C.W. conducted the computational analysis and data interpretation. S.C. annotated the biological data for the AD case study. Y.L., A.M., Q.M., and B.L. wrote the manuscript.

## COMPETING INTERESTS

The authors declare no competing interests.

## Notes

### Competing Interest Statement

The authors have declared no competing interest.

### Summary of Updates

In this revision, the authors re-performed benchmarking analysis and re-generated figures and Supplementary Information.

