## Supplementary Information for "Enhancer-driven gene regulatory networks inference from single-cell RNA-seq and ATAC-seq data"

*To whom correspondence should be addressed

**Supplementary Fig. 1.** A detailed overview of the STREAM framework.

**Supplementary Fig. 2.** Trajectory analyses in MG cells of the AD mouse model brain dataset.

**Supplementary Fig. 3.** Trajectory analyses in AG cells of the AD mouse model brain dataset.

**Supplementary Fig. 4.** Trajectory analyses in OPC cells of the AD mouse model brain dataset.

**Supplementary Fig. 5.** Visualization of an eGRN exclusive to Normal B cells in the DSLL dataset.

**Supplementary Fig. 6.** Visualization of an eGRN exclusive to Tumor B cells in the DSLL dataset.

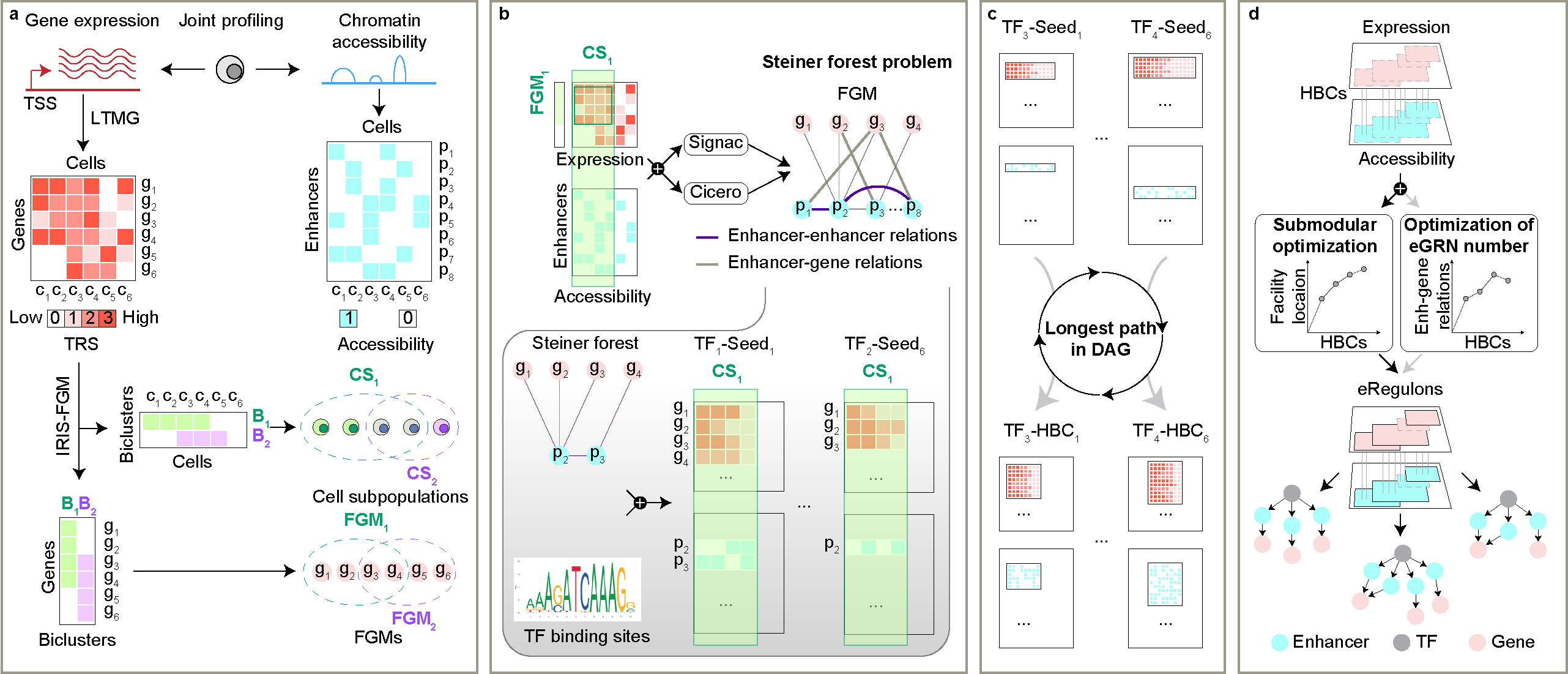

**Supplementary Fig. 1. In-depth schematic of the STREAM framework. a.** Deriving FGMs from joint scRNA-seq and scATAC-seq data profiling. **b.** Using the SFP model for seed element prediction. **c.** Adopting a hybrid biclustering method to discern the longest paths in DAGs. **d.** Applying submodular optimization to determine the optimal eRegulon set.

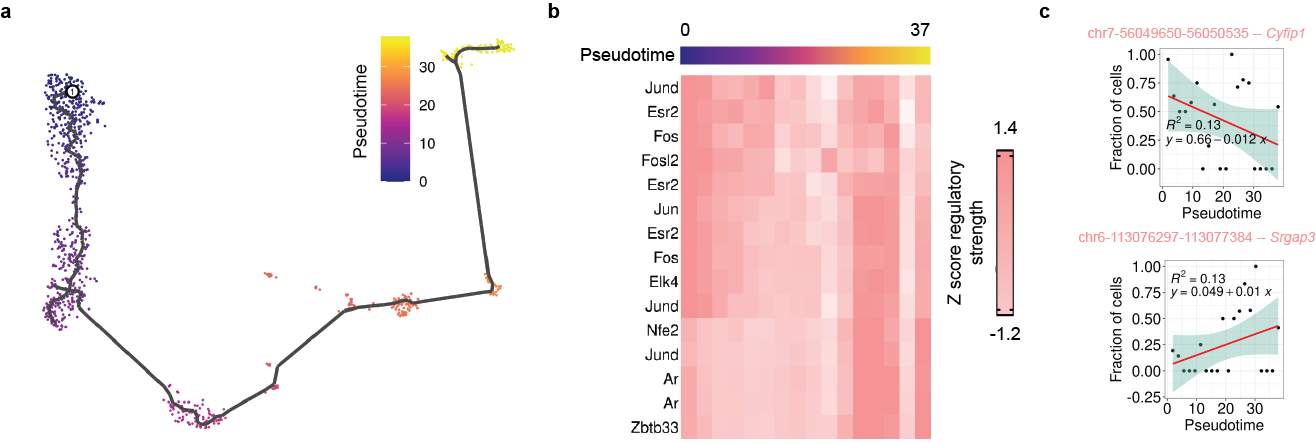

**Supplementary Fig. 2. Trajectory analysis within MG cells from the AD mouse model brain dataset. a.** UMAP visualization color-coded by pseudotime for MG cells. **b.** Heatmap presenting the average regulatory strengths of enhancer-gene relations for MG cell-specific eRegulons across pseudotime. **c-d.** Illustrative enhancer-gene relations exhibiting a consistent trend in regulatory strengths throughout pseudotime in MG cells.

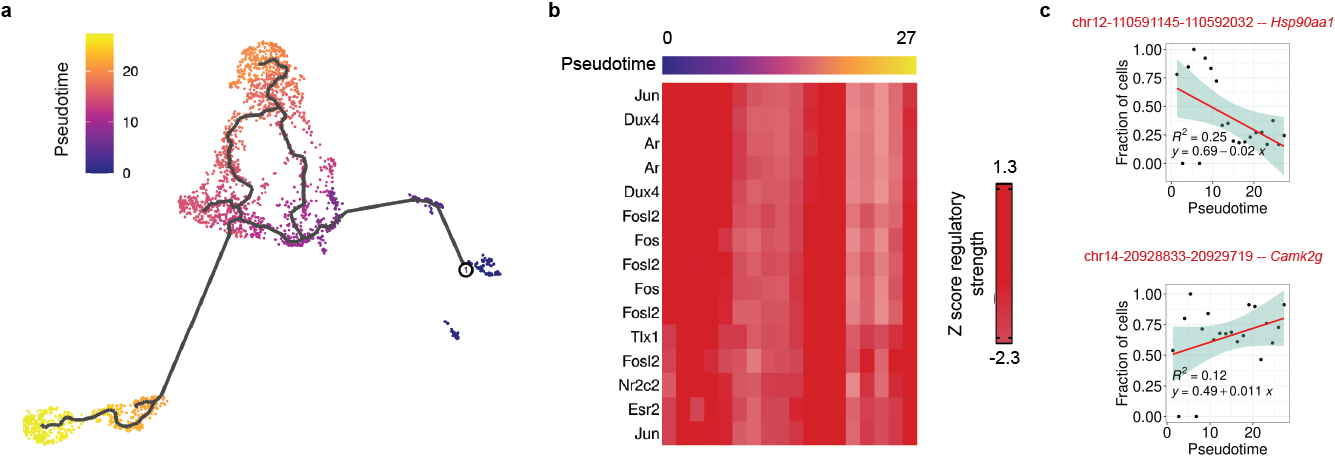

**Supplementary Fig. 3. Trajectory analysis within AG cells from the AD mouse model brain dataset. a.** UMAP visualization color-coded by pseudotime for AG cells. **b.** Heatmap presenting the average regulatory strengths of enhancer-gene relations for AG cell-specific eRegulons across pseudotime. **c-d.** Illustrative enhancer-gene relations showcasing a consistent trend in regulatory strengths throughout pseudotime in AG cells.

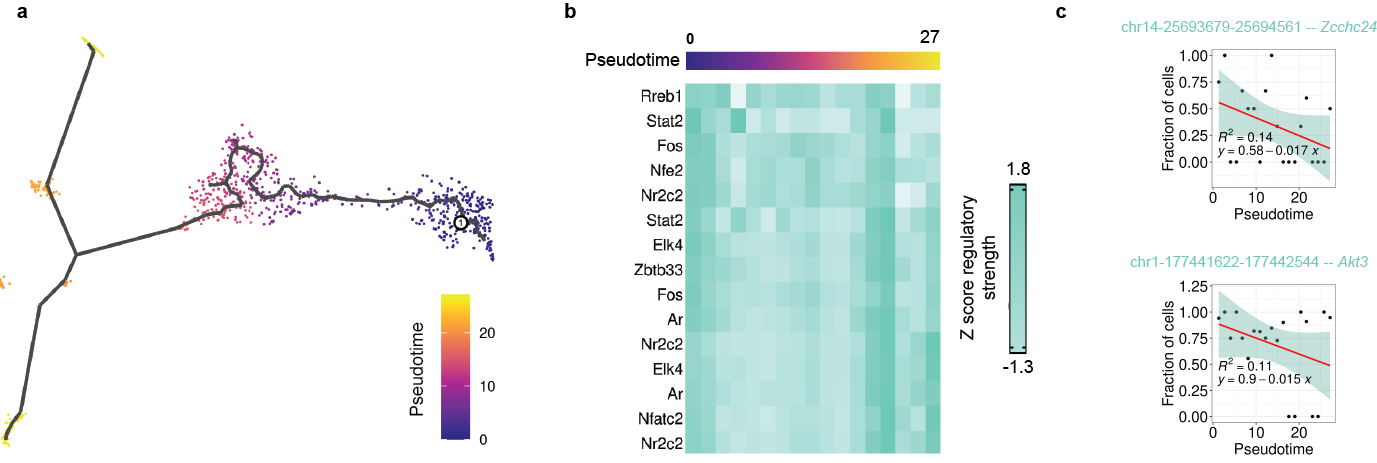

**Supplementary Fig. 4. Trajectory analysis within OPC cells from the AD mouse model brain dataset. a.** UMAP visualization color-coded by pseudotime for OPC cells. **b.** Heatmap illustrating the average regulatory strengths of enhancer-gene relations within eRegulons unique to OPC cells across pseudotime. **c-d.** Representative enhancer-gene relations highlighting a consistent trend in regulatory strengths throughout pseudotime in OPC cells.

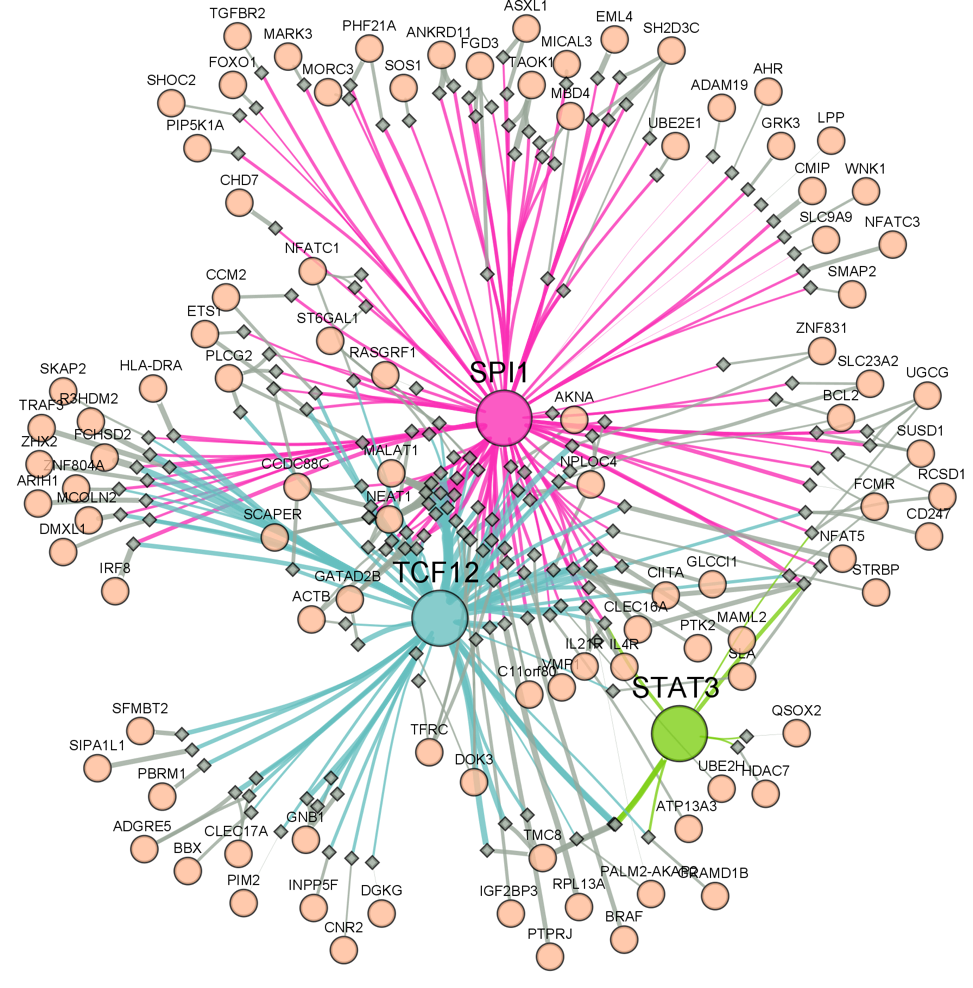

**Supplementary Fig. 5. Depiction of an eGRN unique to Normal B cells within the DSLL dataset.** This eGRN emerges from the synergistic interactions among SPI1, STAT3, and TCF12 and is restricted to particularly variable genes or enhancers. The line thickness between two nodes corresponds to the fraction of cells where both nodes are accessible (for enhancers) or expressed (for genes denoting TFs or their corresponding targets).

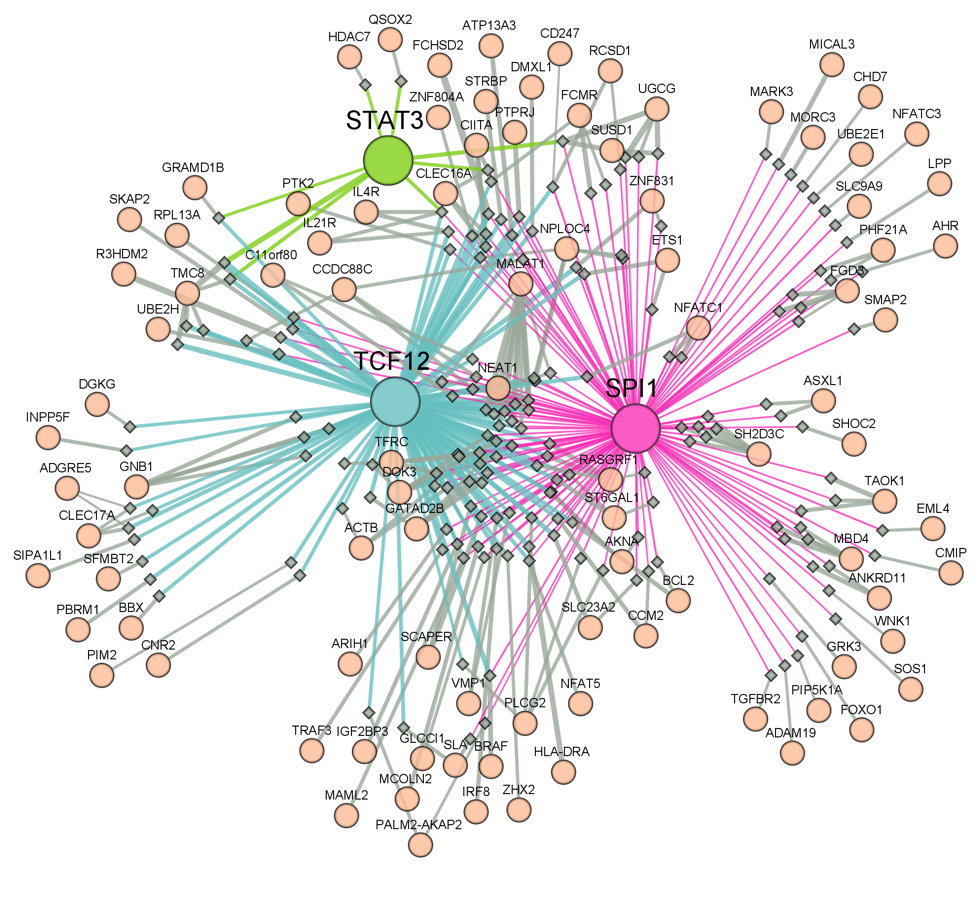

**Supplementary Fig. 6. Depiction of an eGRN unique to Tumor B cells within the DSLL dataset.** This eGRN arises from the collaborative actions of SPI1, STAT3, and TCF12, focusing on notably variable genes or enhancers. The width of the line linking two nodes represents the fraction of cells in which both nodes are either accessible (for enhancers) or expressed (for genes representing TFs or their associated targets).
